# Seeding the meiotic DNA break machinery and initiating recombination on chromosome axes

**DOI:** 10.1101/2023.11.27.568863

**Authors:** Ihsan Dereli, Vladyslav Telychko, Frantzeskos Papanikos, Kavya Raveendran, Jiaqi Xu, Michiel Boekhout, Marcello Stanzione, Benjamin Neuditschko, Naga Sailaja Imjeti, Elizaveta Selezneva, Hasibe Tuncay Erbasi, Sevgican Demir, Teresa Giannattasio, Marc Gentzel, Anastasiia Bondarieva, Michelle Stevense, Marco Barchi, Arp Schnittger, John R. Weir, Franz Herzog, Scott Keeney, Attila Tóth

## Abstract

Programmed DNA double-strand break (DSB) formation is a unique meiotic feature that initiates recombination-mediated linking of homologous chromosomes, thereby enabling chromosome number halving in meiosis. DSBs are generated on chromosome axes by heterooligomeric focal clusters of DSB-factors. Whereas DNA-driven protein condensation is thought to assemble the DSB-machinery, its targeting to chromosome axes is poorly understood. We discovered in mice that efficient biogenesis of DSB-machinery clusters requires seeding by axial IHO1 platforms, which are based on a DBF4-dependent kinase (DDK)–modulated interaction between IHO1 and the chromosomal axis component HORMAD1. IHO1-HORMAD1-mediated seeding of the DSB-machinery on axes ensures sufficiency of DSBs for efficient pairing of homologous chromosomes. Without IHO1-HORMAD1 interaction, residual DSBs depend on ANKRD31, which enhances both the seeding and the growth of DSB-machinery clusters. Thus, recombination initiation is ensured by complementary pathways that differentially support seeding and growth of DSB-machinery clusters, thereby synergistically enabling DSB-machinery condensation on chromosomal axes.

## Introduction

Sexually reproducing eukaryotes employ meiosis to generate haploid reproductive cells from diploid mother cells. One of the key features of meiosis is a specialized homologous recombination that is initiated by programmed formation of DNA double strand breaks (DSBs) at the onset of the first meiotic prophase (reviewed in ^1^). Meiotic DSBs are generated by a type II topoisomerase-related enzyme complex consisting of a catalytic subunit, SPO11, and a co-factor, TOPOVIBL ^2–7^. Meiotic recombination leads to the juxtaposition/synapsis of homologous copies (homologs) of each chromosome in synaptonemal complexes (SCs). Recombination repairs DSBs within the context of synapsed chromosomes, thereby generating inter-homolog genetic exchanges, which produce new allele combinations. Furthermore, a subset of the exchanges mature into chromosomal crossovers (COs), which form the basis of inter-homolog linkages that enable correct chromosome segregation in meiosis. Given the potential genotoxicity of DSBs and the importance of COs, both DSB formation and repair are under tight spatiotemporal control ^8^.

Chromosomes are organized into linear arrays of DNA loops that are anchored on proteinaceous chromosomal cores, called axes, in meiosis ^9^. DSBs are spatially restricted to chromosome axes, which is thought to stem from the concentration of SPO11-activating proteins on axes. SPO11 activity requires several auxiliary proteins in most eukaryotes (reviewed in ^10, 11^). While there is considerable divergence in DSB factors, three SPO11 auxiliary protein families — represented by the budding yeast (*Sc*) Mer2, Mei4 and Rec114 — are conserved in diverse clades of fungi, plants and animals ^12–16^. Both the budding yeast proteins ^17–20^ and the corresponding mouse (*Mm*) proteins (IHO1 (Mer2 ortholog ^14^), MEI4 and REC114) jointly form axis-bound focal clusters (hereafter DSB-factor clusters) that are hypothesized to enable SPO11 activity ^13, 21, 22^. Mammalian DSB-factor clusters incorporate at least two further components, ANKRD31 ^23, 24^, which seems specific to vertebrates, and MEI1 ^25^, orthologs of which are currently known in vertebrates and plants ^26, 27^. *In vitro* studies suggest that DSB factors form chromatin-bound clusters by DNA-driven protein condensation which relies on multivalent protein-protein and protein-DNA interactions ^28^. However, the mechanisms targeting DSB-factor clustering to chromosome axes *in vivo* are not clear.

Axial accumulation of the DSB-machinery was proposed to partially depend on interactions between Mer2/IHO1-family proteins and conserved axis-associated HORMA domain proteins in diverse taxa including fungi ^20, 29, 30^, plants ^31^ and mammals ^22^. In mammals, the HORMA domain protein HORMAD1 is hypothesized to enhance DSB activity ^32, 33^ by enabling the formation of extended axial IHO1 platforms, which serve as substructures for focal clusters of SPO11 auxiliary proteins^22^. Consistent with this hypothesis, (1) axial IHO1 depends on HORMAD1, but not on SPO11 auxiliary proteins ^21, 22^, (2) axial IHO1 accumulations extend beyond the boundaries of focal DSB-factor clusters ^22^, (3) focal DSB-factor clusters largely depend on IHO1 and, to a lower extent, HORMAD1 in most of the genome ^22, 24^, and (4) DSBs depend fully on IHO1 ^22^ and partly on HORMAD1 ^32, 33^. In line with these observations, HORMAD1 regulation is thought to enable correct spatiotemporal patterning of DSB activity. In several studied models, SC seems to limit DSB-machinery ^13, 18, 20, 22, 31, 34–36^ and DSB activity ^8, 37–39^ to unsynapsed sections of axes where DSBs are used for homolog pairing and synapsis. This regulatory mechanism was hypothesized to involve an SC-triggered depletion of HORMAD1 from synapsed axes in mammals ^40^.

DSBs have alternative requirements for SPO11 auxiliary proteins in the relatively short (∼0.7 Mb in mice) pseudoautosomal regions (PARs) of sex chromosomes, where DSB-factor levels ^23–25^ and DSB activity ^41^ are greatly elevated to enable X and Y chromosome pairing in males. Whereas ANKRD31 is not essential for DSB–factor clusters and DSBs in most of the genome, ANKRD31 is critical in the PAR ^23, 24^. Neither IHO1 ^24^ nor HORMAD1 ^25^ is needed for enrichment of SPO11 auxiliary proteins on PAR axes.

Whereas the dependency of DSB-machinery clusters on HORMAD1 and IHO1 has been extensively studied (see previous paragraphs), it remains unknown if and how the HORMAD1-IHO1 interaction enables assembly of DSB-machinery on axes. Here, we reveal the mechanism that the C-terminal 7 amino acids of IHO1 are required for (1) IHO1-HORMAD1 interaction, (2) the formation of axial IHO1 platforms and (3) efficient seeding of cytologically distinguishable DSB-factor clusters. These observations collectively suggest that seeding of the DSB machinery on axes critically complements and enhances the previously suggested mechanism of DSB-factor clustering by DNA-driven condensation *in vivo*. We also discover that whereas IHO1-HORMAD1 interaction specifically enhances seeding of DSB-factor clusters, ANKRD31 supports both seeding and growth. The IHO1-HORMAD1 complex and ANKRD31 act synergistically — their simultaneous disruption abolishes DSB-factor clusters and DSBs. Thus, DSB formation is enabled on chromosome axes by complementary pathways that differ in both mechanism and preferred genomic locations.

## Results

### IHO1-HORMAD1 interaction requires a conserved acidic motif in the IHO1 C-terminus

A direct HORMAD1-IHO1 interaction may enable focusing of DSB activity to unsynapsed axes^22^. Therefore, we mapped HORMAD1-interacting regions of IHO1 in yeast two-hybrid (Y2H) assays (Fig 1A-B, Table S1). The first 358 (of 574) amino acids of IHO1, including a conserved coil domain, were neither sufficient nor required for interaction with HORMAD1. In contrast, IHO1 fragments that included the C-terminal 75 amino acids of IHO1 interacted with HORMAD1 and, specifically, its HORMA domain. Further, *in vitro* binding assays reconfirmed efficient interaction between the HORMA domain of HORMAD1 and the C-terminal 215 amino acids of IHO1 (Fig. S1A-B).

**Figure 1.**
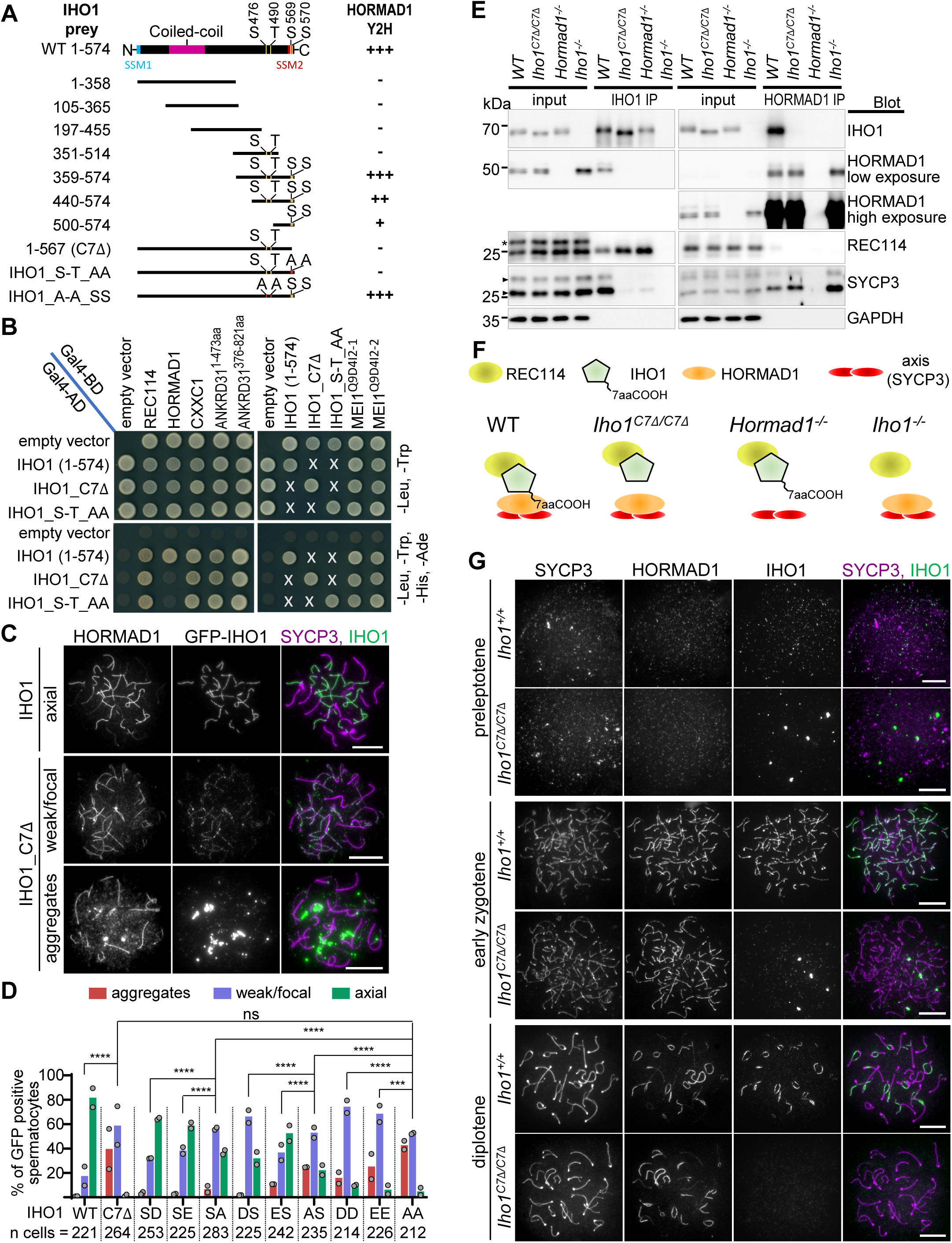
IHO1 C-terminus mediates IHO1-HORMAD1 interaction **A-B** Y2H assays between IHO1 interactors and wild-type (1-574) or modified versions of IHO1. **A** Schematics show conserved domains ^14^ and positions of phospho-serines (S) or -threonine (T) or their substitution with alanine (A) in IHO1. **B** Yeast cultures are shown after 3 (left) or 2 (right) days of growth. **C, G** Immunostaining in nuclear spread spermatocytes of 13 days postpartum (dpp) (**C**) and adult (**G**) mice. Chromosome axis (SYCP3, **C overlay, G)**, HORMAD1 (**C**, **G**) and either ectopically expressed GFP-IHO1 (**C**) or endogenous IHO1 (**G**) were detected. Bars, 10 µm. **D** Quantification of localization of GFP-tagged IHO1 versions in late zygotene. IHO1 versions: wild type (WT), a mutant missing the last 7 amino acids (C7Δ) and versions where single-letter amino acid code indicates point mutations in positions 569 and 570. Block bars are means. Likelihood-ratio test, ns=P>0.05, ***=P<0.001, ****=P<0.0001. **E** Immunoprecipitation (IP) immunoblots from testis extracts of 13 dpp mice. Asterisk and triangles mark unspecific protein band in REC114 blot and isoforms of SYCP3, respectively. **F** Schematics summarizing conclusions of panel E. See also Figure S1, Table S1 and S2.

IHO1 is phosphorylated *in vivo* on at least four positions, S476, T490, S569 and S570 (^42^, see https://phosphomouse.hms.harvard.edu/site_view.php?ref=IPI00914141), which are also located in the C-terminal region of IHO1 (Fig 1A). Simultaneous replacement of S476 and T490 of IHO1 by alanine did not affect the Y2H interaction of IHO1 and HORMAD1. In contrast, IHO1-HORMAD1 interactions were abolished by (i) simultaneous exchange of serines to alanines in positions 569 and 570 (Fig. 1A-B) or (ii) the deletion of the last 7 amino acids of IHO1 (hereafter IHO1_C7Δ, Fig. 1A-B, S1A-B, Table S1). All known IHO1 interactors except HORMAD1 efficiently interacted with IHO1_C7Δ (Fig. 1B, Table S1), supporting the idea that the C-terminus of IHO1 is specifically important for interaction with HORMAD1 (via the HORMA domain).

S569 and S570 of IHO1 are located in an acidic FDS_(569)_S_(570)_DDD sequence patch that overlaps with the C-terminal end of a widely conserved short similarity motif (called SSM2) of Mer2/IHO1-family proteins ^14^ (Fig. 1A, S1C). Clusters of acidic residues are almost universally present in or next to SSM2s, and one or more serine/threonine(s) are often found in the acidic patch, particularly in vertebrates and plants (^14^, Fig. S1C, Table S2).

Consistent with their conservation, the C-termini of Mer2/IHO1-family proteins are important for interactions with HORMAD1 orthologs in several taxa, including budding ^29^ and fission ^30^ yeasts in addition to mammals (Table S2). IHO1 and HORMAD1 orthologs also interact in plants ^31, 36, 43, 44^. In *Arabidopsis thaliana* (*At*), the coiled coil-containing N-terminus of *At*Mer2/IHO1 (PRD3) — but not the SSM2-harboring C-terminus — was reported to interact with the SWIRM domain-harboring C-terminus of the *At*HORMAD1 (ASY1) ^36^. In contrast, we found that the C-terminus of *At*PRD3 interacted with the HORMA domain-containing N-terminus of *At*ASY1 in low stringency Y2H, and that the interaction required the SSM2-linked acidic patch of *At*PRD3 (Fig. S1D). Thus, conserved acidic patches associated with SSM2s may enable and/or enhance interaction of IHO1-and HORMAD1-related proteins in diverse taxa, albeit the importance and the molecular role of SSM2/acidic patches may differ between species (summarized in Table S2).

### IHO1-axis association requires the IHO1 C-terminus

To test if the C-terminal region of IHO1 was important for IHO1 localization to chromosomes we ectopically expressed GFP fusions of wild type and mutant versions of IHO1 in spermatocytes by *in vivo* electroporation of mouse testes (Fig. 1C-D). IHO1 mutations included a deletion of the last 7 amino acids, or an exchange of S569 and/or S570 for either non-phosphorylatable alanine, or phosphomimetic aspartates or glutamates. All of the tested mutations impaired axial localization of the GFP-IHO1 fusions, but serine to alanine mutations resulted in more severe defects than serine to aspartate or glutamate exchanges in respective positions. IHO1 localization was most severely disrupted if either the last 7 amino acids were deleted or both S569 and S570 were exchanged with alanines. These versions of IHO1 rarely accumulated effectively on axes, although weak focal staining on chromatin and/or aggregate formation was often observed. These observations suggest that the IHO1 C-terminus enhances axial recruitment of IHO1 by promoting IHO1-HORMAD1 interaction. Whereas S569 and S570 seem to have a critical function, we are cautious with interpreting the phenotypes caused by non-phosphorylatable and phosphomimetic replacements at these sites. Hence, the significance of *in vivo* phosphorylation at S569 and S570 remains uncertain.

To test if disruption of the C-terminus also altered the behavior of endogenous IHO1 we generated a mutant mouse line expressing IHO1_C7Δ from a gene-edited *Iho1* locus (*Iho1^C7Δ/C7Δ^* genotype, Fig S1E). Whereas deletion of the last 7 amino acids of IHO1 did not significantly reduce testicular IHO1 levels (Fig. 1E, Fig. S1F), it changed the protein interactions of IHO1, as assayed by immunoprecipitation in testes extracts (Fig. 1E-F, Fig. S1G). Wild-type IHO1 formed complexes with HORMAD1, the core axis component SYCP3 and the DSB-factor REC114. In contrast, IHO1_C7Δ efficiently formed complexes with REC114, but not with HORMAD1 or SYCP3 in *Iho1^C7Δ/C7Δ^* mice, which paralleled loss and persistence of IHO1-SYCP3 and IHO1-REC114 complexes, respectively, in *Hormad1^-/-^* mice. These results indicate that the C-terminus of IHO1 enables IHO1-HORMAD1 complex formation, thereby promoting a link between the DSB-machinery and the meiotic chromosome axis.

Importantly, the loss of either HORMAD1 or the last 7 amino acids of IHO1 caused a depletion of IHO1 from chromatin-enriched fractions of testis extracts (Fig. S1H). Further, resembling localization of wild-type IHO1 in *Hormad1^-/-^* meiocytes, IHO1_C7Δ did not efficiently accumulate on unsynapsed chromosome axes in either sex of HORMAD1-proficient meiocytes (Fig. 1G, Fig. S1I-J). Instead, IHO1_C7Δ formed aggregates, which associated with a few (typically 3-4) chromosome ends in a late zygotene-like stage (Fig. S1I). Together, our observations suggest that loss of the last 7 amino acids prevents endogenous IHO1 from efficiently binding HORMAD1 and axes without significantly affecting its other known protein interactions.

In fission yeast, deletion of the C-terminal SSM2/acidic patch of the Mer2/IHO1 ortholog analogously disrupted its interactions with orthologs of both HORMAD1 and the core axis components SYCP2/SYCP3, resulting in its loss from chromosome axes ^30^. Thus, the C-terminal SSM2/acidic patch promotes the axis association of Mer2/IHO1-family proteins in distantly related species.

### DDK-dependent phosphorylation of the IHO1 C-terminus enhances axial accumulation

SDS-PAGE revealed a fast and a slow migrating form of IHO1 (Fig. 2A-C, S2A) ^22^. The slow-migrating form was more abundant in chromatin-enriched fractions of testis extracts of wild type (Fig. 2A, S1H), suggesting that IHO1 was posttranslationally modified in correlation with chromatin binding. The slow-migrating form was present in *Hormad1^-/-^* mice (Fig. 2B), where IHO1 was depleted from chromatin (Fig. S1H) ^22^. Hence, IHO1 modifications may be a cause rather than an outcome of IHO1 chromatin binding. The slow-migrating form was also present in *Spo11^-/-^* mice indicating that it did not require DSB formation (Fig. 2B, S1H). Phosphatase treatment converted the slow-into the fast-migrating form (Fig. 2C, S2A), indicating that the slow-migrating form represented phosphorylated IHO1. IHO1_C7Δ lacks two of the four known phospho-sites of wild-type IHO1. Immunoblot analysis detected only a single IHO1_C7Δ protein band, which had a slightly higher electrophoretic mobility than the fast-migrating form of wild-type IHO1 (Fig. 2A-C). Thus, loss of the last 7 amino acids, and the S569/570 phospho-sites within, may have prevented phosphorylation of all sites in IHO1. Alternatively, in the absence of S569/570, phosphorylation of S476 and T490 or other unknown sites were insufficient for a discernible mobility shift in standard SDS-PAGE. The latter hypothesis was supported by analysis on phos-tag gels (Fig. S2B), which enable detection of distinct phosphoforms of proteins by enhancing their retardation during electrophoresis ^45, 46^.

**Figure 2.**
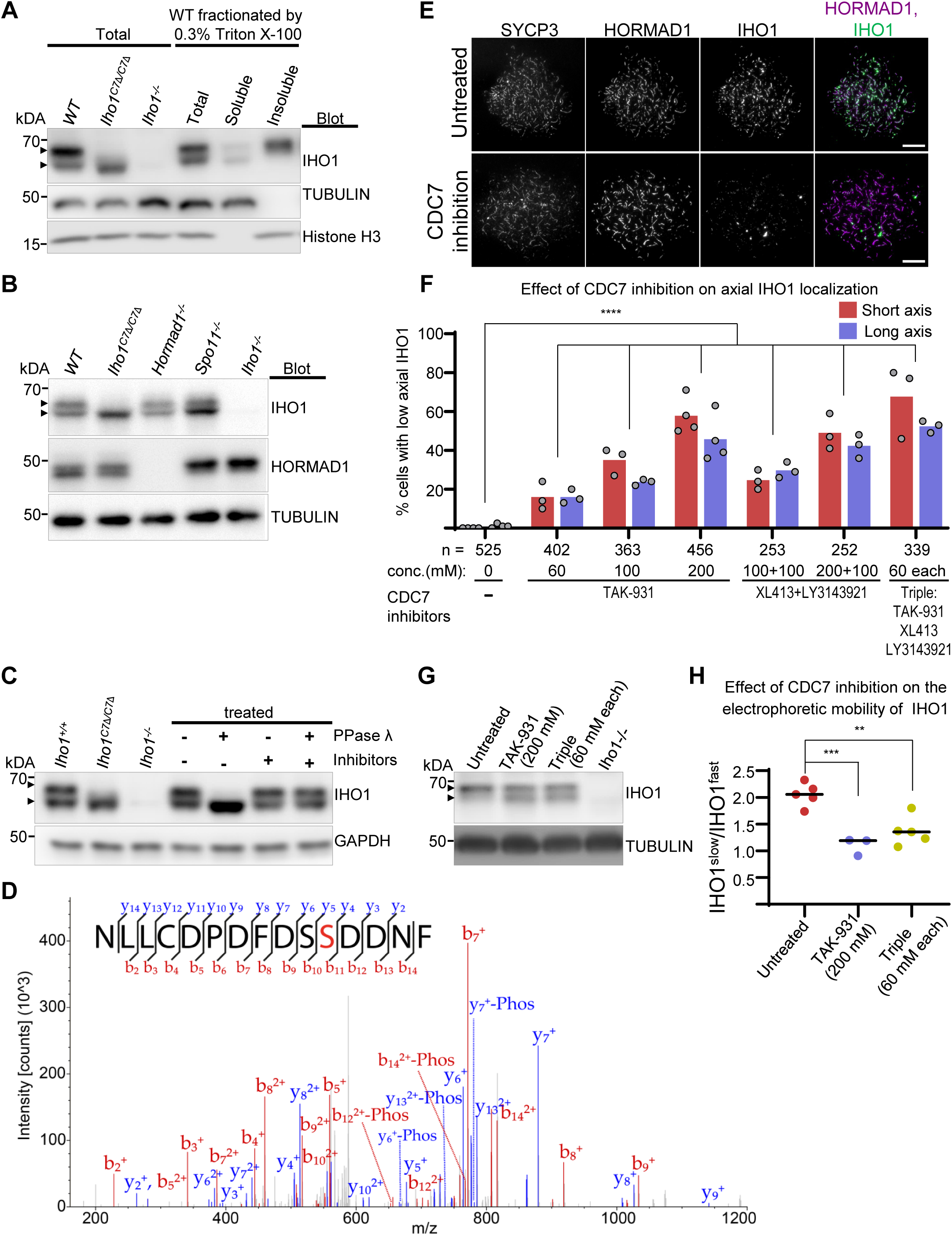
CDC7-dependent phosphoform of IHO1 is enriched in chromatin **A-C** SDS-PAGE immunoblots of protein extracts from testes of 13 dpp mice. Total (**A-C**), fractionated (**A**) and/or phosphatase treated total (**C**) extracts are shown. **D** MS/MS spectrum of NLLCDPDFDS(pS)DNF-COOH peptide. Identified b and y ions are annotated in red and blue, respectively. Fragments with neutral loss of H_3_PO_4_ are indicated as “-Phos”. 10.9+/-1.4% of the peptide was phosphorylated in 2 measurements. **E-H** Analysis of testes of 8 dpp mice after 48 hours *in vitro* culture with or without CDC7 inhibitors. **E** Nuclear spread spermatocytes. Bars, 10 µm. **F** Quantification of spermatocytes with depleted IHO1 on axes. Block bars are means. Likelihood ratio test, ****=P< 2.20E-16. **G** SDS-PAGE immunoblots of testis extracts. **A-C, G** Arrowheads mark slow and fast migrating IHO1 forms; slow migrating forms were reproducibly more dominant in untreated testis cultures (**G**) than in freshly processed testes (**A-C**). **H** Intensities of slow migrating IHO1 band normalized to intensities of fast migrating bands from SDS-PAGE immunoblots. Bars are means. Two tailed *t* test, **=P< 0.01, ***=P<0.001. **G-H** “Triple” inhibitor mix, as in **F**. See also Figure S2, Table S3.

S570 is followed by acidic aspartates, characteristic of potential target sites of the DBF4-dependent CDC7 kinase (DDK) ^47, 48^. DDK is a key activator of the budding yeast IHO1 ortholog, Mer2 ^49–51^, hence DDK may regulate IHO1 in mice. DBF4 interacted with both IHO1 and IHO1_C7Δ in Y2H (Fig. S2C). Further, recombinant DDK phosphorylated peptides corresponding to the last 15 amino acids of IHO1 in an *in vitro* kinase assay; phosphorylation was detected on one but not both serines in about 10% of peptides by mass spectrometry (Fig. 2, S2D and Table S3). The serine that corresponded to S570 of IHO1 was identified as a phospho site with a probability of more than 99.6% by about 50% of annotated spectra. The remaining spectra were annotated with a site probability of 50% for S569 and S570, indicating that only phosphorylation on S570 has been unambiguously identified. These observations indicate that DDK inefficiently phosphorylates S569 *in vitro*, but DDK may phosphorylate both S569 and S570 in the context of full length IHO1 *in vivo*.

To address if DDK regulated IHO1 in meiocytes we exposed testis organ cultures to distinct DDK inhibitor cocktails. DDK inhibition impaired IHO1 localization to axes (Fig. 2E-F) and reduced the abundance of the slow-migrating phospho-band of IHO1 in SDS-PAGE (Fig. 2G-H). Further, DDK inhibition lowered recombination focus numbers, as detected by DMC1 staining, in both wild-type and *Iho1^C7Δ/C7Δ^* spermatocytes (Fig. S2E-G). These observations suggest that DDK promotes recombination through phosphorylation of the IHO1 C-terminus — likely enhancing both axial IHO1 levels and DSB activity — but also independently of the IHO1 C-terminus. In contrast to the role of DDK in mice, DDK-mediated phosphorylation of the budding yeast Mer2 is required for DSB activity but not axial recruitment of Mer2 ^20, 49–51^. Thus, reliance of recombination initiation on DDK is conserved between mice and yeast, with a divergence in the nature of DDK-controlled functions.

### Elevated loss of meiocytes in *Iho1^C7Δ/C7Δ^* mice

*Iho1^C7Δ/C7Δ^* male mice did not display infertility (Table S4), but they had 1.45-fold smaller testis size (Fig. 3A), and an elevated loss of spermatocytes from mid pachytene stage until meiotic divisions (Fig. S3A-B, Table S5). Female *Iho1^C7Δ/C7Δ^* mice were also fertile (Table S4), but oocyte numbers were approx. 2-fold lower in young *Iho1^C7Δ/C7Δ^* than wild type mice (Fig. S3C-D), and fertility of *Iho1^C7Δ/C7Δ^*females declined prematurely (Table S4). Meiotic recombination defects trigger elimination of both spermatocytes and oocytes ^52–55^, therefore *Iho1^C7Δ/C7Δ^* phenotypes were consistent with disrupted meiotic recombination. The phenotypes were milder in *Iho1^C7Δ/C7Δ^*mice as compared to *Hormad1^-/-^* mice, where inefficient axial recruitment of IHO1 was accompanied by (i) infertility, (ii) a complete elimination of spermatocytes at a mid pachytene-like stage and (iii) a >3-fold lower testis size as compared to wild type (Fig. 3A) ^32, 33,56^. Hence, despite similar depletion of IHO1 from axes, meiotic recombination defects are less severe in *Iho1^C7Δ/C7Δ^* than *Hormad1^-/-^*.

**Figure 3.**
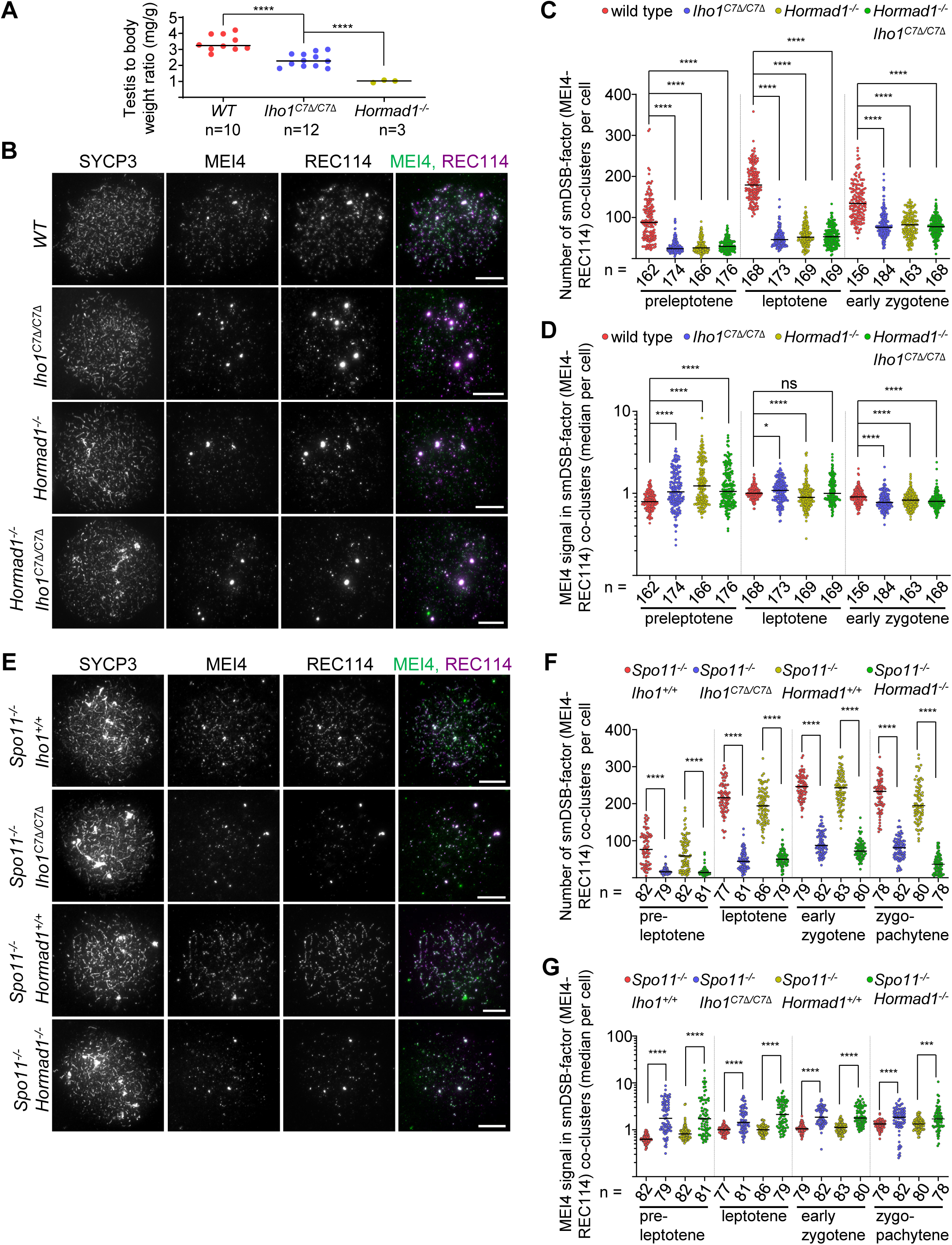
*Iho1^C7Δ/C7Δ^* mice show reduction in testis size and DSB-factor cluster numbers **A** Testis to body weight ratios in adult mice (age 50-120 days). Bars mark means. Two tailed Welch t-test, s****=P<0.0001. **B, E** Immunostaining in nuclear spread leptotene spermatocytes of adult mice. Bars, 10 µm. **C, D, F, G** Numbers of small MEI4-REC114 co-clusters (**B, E**) and MEI4 intensities in MEI4-REC114 co-clusters (**C, F**, data points show median cluster intensities per cell) in spermatocytes of adult mice. Zygo-pachytene (**F-G**) is equivalent to a mix of late-zygotene and early pachytene stages which are indistinguishable in SC-defective backgrounds. Bars are medians, n=cell numbers. Mann Whitney U-Test, ns=P>0.05, *=P< 0.05, ***=P<0.001, ****=P<0.0001. See also Figure S3, Table S4 and S5.

### Diminished presence of DSB-factors on chromosome axes in *Iho1^C7Δ/C7Δ^* meiocytes

To further assess the role of the IHO1 C-terminus in recombination, we compared deletions of the IHO1 C-terminus, HORMAD1, or both, for their impact on DSB-factor clusters in spermatocytes (Fig. 3 and S3E-F). DSB-factor clusters exist in two types — small clusters (hereafter smDSB-factor clusters) on axes across the genome and large clusters (hereafter laDSB-factor clusters) in mo-2 minisatellite-rich regions, which include PARs and PAR-like telomeric regions of three autosomes ^23, 24^. IHO1 and HORMAD1 are important for smDSB-factor clusters but dispensable for laDSB-factor clusters ^22, 24, 25, 57^. Consistent with the known functions of IHO1 ^22, 24^, we observed efficient assembly of laDSB-factor clusters (as represented by large, >1 µm diameter MEI4 foci) at PARs and a subset of autosomal ends in both wild-type and *Iho1^C7Δ/C7Δ^*spermatocytes (Fig. S3E-F). In contrast, numbers of smDSB-factor clusters (detected as small REC114-MEI4 co-foci) were strongly reduced (1.8 to 3.9 fold) in *Iho1^C7Δ/C7Δ^* spermatocytes as compared to wild type (Fig. 3B-C). Numbers of smDSB-factor clusters were reduced to similar extents in *Iho1^C7Δ/C7Δ^*, *Hormad1^-/-^* and *Hormad1^-/-^ Iho1^C7Δ/C7Δ^* spermatocytes (<1.2-fold differences at matched stages, Fig. 3B-C), suggesting that deletion of IHO1 C-terminus and loss of HORMAD1 impair the same pathway of DSB-machinery assembly.

Although the numbers of smDSB-factor clusters were reduced throughout early prophase, their median signal intensities were significantly higher in *Iho1^C7Δ/C7Δ^*, *Hormad1^-/-^* and *Hormad1^-/-^ Iho1^C7Δ/C7Δ^*than in wild-type spermatocytes in the preleptotene stage, when DSBs have not yet formed (Fig. 3D). These signal intensities then gradually dropped below wild-type levels in all three mutants upon progression to early zygotene, when early recombination intermediates reach their peak levels and DSB activity is thought to diminish in wild type. Seeing fewer but brighter clusters at early stages suggests that the seeding of cytologically distinguishable DSB-factor clusters is defective but their initial growth is efficient in the absence of IHO1 C-terminus and/or HORMAD1. DSBs and downstream DNA damage signaling disrupt the DSB-machinery by multiple pathways ^34^. Therefore, progressive diminution of smDSB-factor clusters in *Iho1^C7Δ/C7Δ^*, *Hormad1^-/-^* and *Hormad1^-/-^ Iho1^C7Δ/C7Δ^* spermatocytes by zygotene might indicate that smDSB-factor clusters are prone to enhanced destabilization by DSBs if IHO1 and HORMAD1 functions are impaired.

To test this hypothesis, we assessed how deletion of the IHO1 C-terminus or absence of HORMAD1 affected smDSB-factor clusters in *Spo11^-/-^*spermatocytes, where programmed DSBs do not form (Fig. 3E-G). SmDSB-factor clusters were fewer in number but had consistently higher median signal intensities in *Iho1^C7Δ/C7Δ^ Spo11^-/-^* and *Hormad1^-/-^ Spo11^-/-^* spermatocytes as compared to *Spo11^-/-^* spermatocytes in all prophase stages that were present in these genotypes. These observations support the conclusion that smDSB-factor clusters require HORMAD1 and the IHO1 C-terminus for both efficient seeding and resistance to destabilization by DSBs, but not growth *per se*.

### The IHO1 C-terminus is required for efficient DSB formation

Given the depletion of smDSB-factor clusters in the absence of the IHO1 C-terminus, we tested if DSB formation was also reduced. Processing of SPO11-generated DSBs results in SPO11-oligo complexes, whose testicular levels inform about DSB activity, allowing comparison of DSB formation in conditions where cellular compositions of testes are similar ^58^. Meiotic recombination defects trigger apoptosis in mid pachytene, which is reached at 14 days postpartum (dpp) in the first wave of meiosis in our strain background. Hence, testis cellularity is unaltered by meiotic recombination defects before 14 dpp. Testicular levels of SPO11-oligo complexes were reduced in both *Iho1^C7Δ/C7Δ^* and *Hormad1^-/-^* as compared to wild type mice at 13 dpp, which indicated that both the IHO1 C-terminus and HORMAD1 were needed for efficient DSB formation (Fig. 4A-B). Deletion of the IHO1 C-terminus did not cause a further reduction of SPO11-oligo levels in a *Hormad1^-/-^* background (Fig. S4A). This supports the hypothesis that loss of the IHO1 C-terminus principally disrupted only those functions of IHO1 in DSB formation that are HORMAD1 dependent.

**Figure 4.**
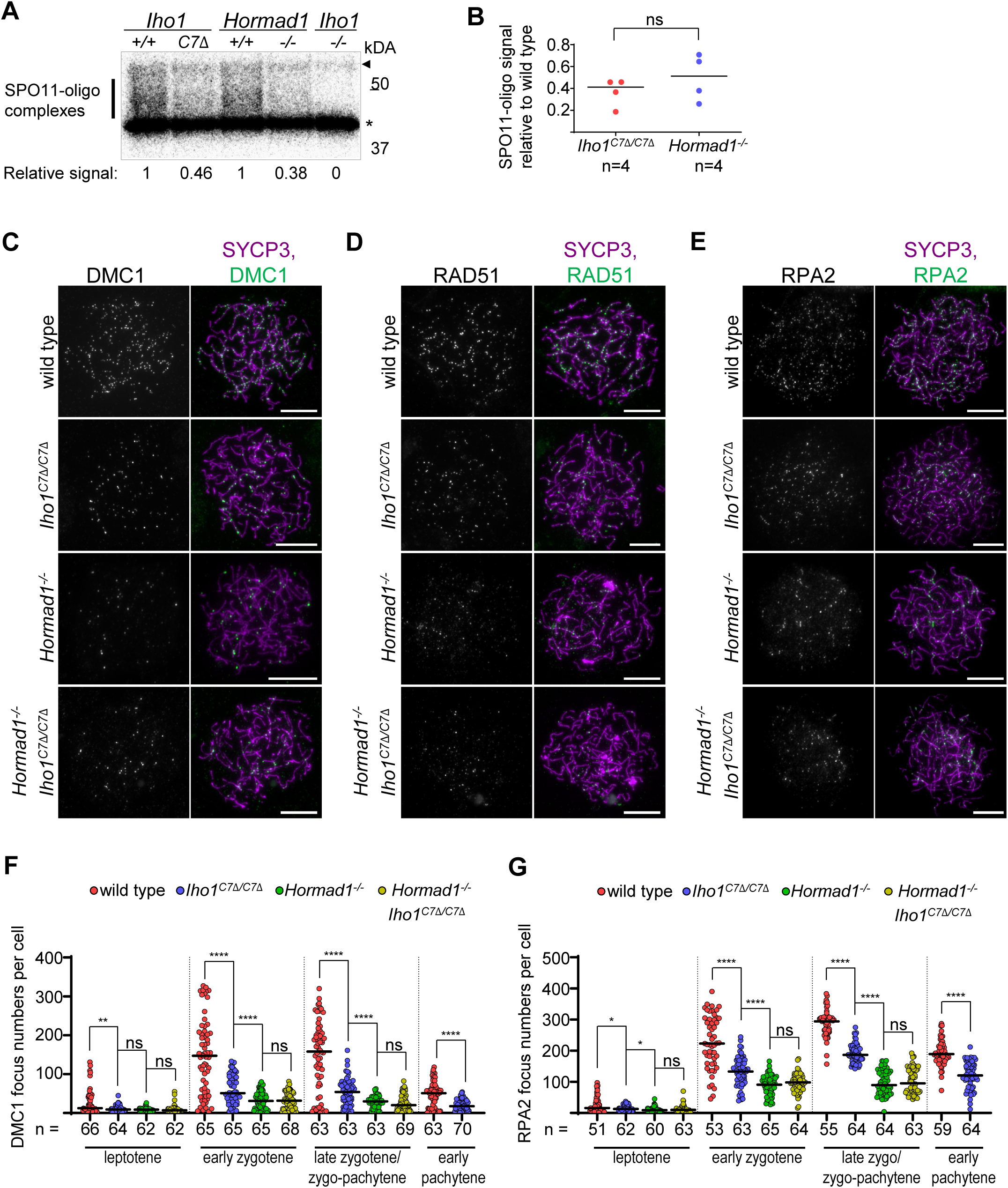
Efficient meiotic recombination relies on IHO1-HORMAD1 interaction **A** Radiograph of immunoprecipitated and radioactively labeled SPO11-oligo complexes from testes of 13 dpp juvenile mice. Bar, SPO11-specific signals, asterisk, nonspecific labelling, and arrowhead, immunoglobulin heavy-chain. Radioactive signals were background-corrected (*Iho1^-/-^* , signal=0) and normalized to corresponding wild type control (1). **B** Quantification of SPO11-oligo complexes from 13dpp mice. Bars are mean, n=number of mice. Paired t-test, ns=P>0.05. **C-E** Immunostaining in nuclear spread early zygotene spermatocytes of adult mice. Bars, 10 µm. **F-G** Quantification of axis associated DMC1 (**F**), RPA2 (**G**) focus numbers in spermatocytes. Bars are medians, n=cell numbers. Mann Whitney U-test, ns=P>0.05, *=P< 0.05, **=P<0.01, ****=P<0.0001. See also Figure S4.

Processing of DSBs generates single-stranded DNA ends that initiate recombination by invading homologous DNA sequences with the help of recombination proteins that accumulate — manifesting as foci — on single-stranded DNA ends. To further compare the roles of HORMAD1 and IHO1 C-terminus in recombination initiation we examined foci of recombination proteins, DMC1, RAD51 and RPA2, in meiocytes (Fig. 4C-G and S4B-H). *Iho1^C7Δ/C7Δ^* mice showed significantly reduced recombination focus numbers in both spermatocytes (medians, 1.6-2.9 fold in zygotene, Fig. 4F-G, S4B) and oocytes (medians, 2.3-3.1 fold in zygotene, Fig. S4F-H) as compared to wild type. Further markers of DSBs, phosphorylated histone H2AX (ɣH2AX) flares on synapsed autosomes, were also diminished in pachytene stage *Iho1^C7Δ/C7Δ^* spermatocytes as compared to wild type (Fig. S4I-J). Together, these observations indicated that DSB activity was reduced in *Iho1^C7Δ/C7Δ^* mice, which likely accounted for defects in meiotic progression and/or fertility.

Interestingly, whereas the low recombination focus numbers were similar in *Hormad1^-/-^* and *Hormad1^-/-^ Iho1^C7Δ/C7Δ^*spermatocytes, focus numbers were consistently lower in these genotypes than in *Iho1^C7Δ/C7Δ^* single mutants (Fig. 4C-G and S4B). The former observation reconfirms that loss of IHO1 C-terminus impairs only HORMAD1-dependent functions of IHO1 in DSB formation. The latter observation suggests that HORMAD1 modulates recombination beyond recruiting IHO1 to chromosome axes. This conclusion is in agreement with work in budding yeast, where the HORMAD1 ortholog Hop1 not only promotes DSBs, but also enables normal DSB repair kinetics and enhances the use of homologs instead of sister chromatids as recombination partners ^59–61^. Previously reported *Hormad1^-/-^* phenotypes were also consistent with a HORMAD1 role in DSB repair ^62, 63^, and HORMAD1 was hypothesized to prevent premature turnover of early recombination intermediates, thereby sustaining high levels of single-stranded DNA ends for homology search ^40, 63^. Thus, a conserved HORMAD1 function impacting DSB repair kinetics and/or template choice may explain more severe recombination defects in *Hormad1^-/-^*than *Iho1^C7Δ/C7Δ^* spermatocytes.

### Assurance of synapsis and CO formation between homologs requires the IHO1 C-terminus

DSBs are needed in high numbers to enable efficient synapsis and crossover formation between each homolog pair in mice ^39^. Synapsis is not completed in most spermatocytes if DSB activity is below 50% of wild-type levels according to analysis of a *Spo11*-hypomorphic mutant mouse line, *Tg(Spo11β)^+/-^* ^39^. Approximately twofold fewer DSBs were found in *Iho1^C7Δ/C7Δ^* as compared to wild type (Fig. 4 and S4A-H), hence we tested if reduced DSB activity resulted in downstream synapsis errors in *Iho1^C7Δ/C7Δ^* meiocytes.

A subset of *Iho1^C7Δ/C7Δ^* meiocytes formed synaptonemal complexes (SCs) on all chromosomes, but efficiency and accuracy of SC formation was reduced in *Iho1^C7Δ/C7Δ^* as compared to wild type in both sexes (Fig. 5A-D Fig. S5A-E). SCs connected PARs of sex chromosomes and the full length of autosomes in most wild-type spermatocytes in pachytene stage. In contrast, pachytene-like *Iho1^C7Δ/C7Δ^* spermatocytes featured four main types of synapsis defects, (i) non-homologous entanglements/synapsis, (ii) partially unsynapsed autosomes, (iii) fully unsynapsed PARs and/or (iv) autosomes (Fig. 5B-C and Fig. S5A). Whereas frequencies of non-homologous synapsis and autosomal asynapsis decreased as pachytene progressed, sex-chromosome asynapsis grew more frequent (Fig. 5B). In most of the asynaptic *Iho1^C7Δ/C7Δ^*spermatocytes (n=175), one to two autosomes were affected (average 1.12).

**Figure 5.**
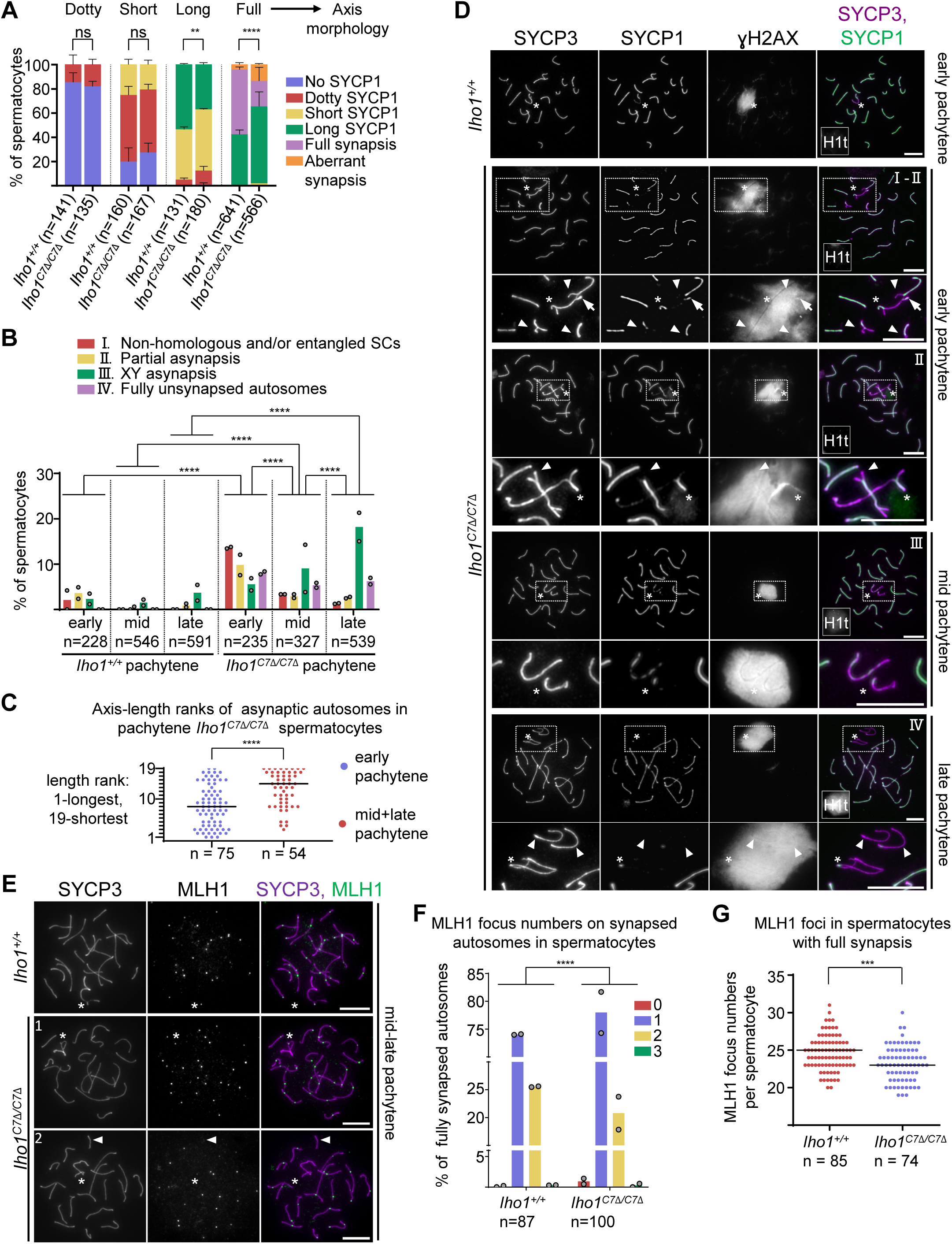
Efficient SC and CO formation require IHO1-HORMAD1 interaction **A-C, F-G** Quantifications of SC morphology relative to axis development (**A**), I-IV non-exclusive SC defect types (**B**), axis-length rank of asynaptic autosomes (**C**), MLH1 focus numbers on synapsed autosomes (**F**) or total MLH1 numbers (**G**) in spermatocytes of adult mice. Leptotene to early pachytene cells (identified by a lack of histone H1t) (**A**), pachytene cells (**B-C**), or both fully synapsed and asynaptic (**F**) or only fully synapsed (**G**) late pachytene cells were examined. Pools of two experiments are shown. Bars are means (**A-B, F**) or medians (**C, G**), error bars are standard deviation (**A**), n=numbers of spermatocytes. Likelihood ratio test (**A-B, F**) or Mann Whitney U-Test (**C, G**), ns=P>0.05, **=P<0.01, ***=P<0.001, ****=P<0.0001. **D-E** Immunostaining in nuclear spread spermatocytes of adult mice. Asterisks, sex chromosomes. **D** Insets are enlarged under respective panels. Arrowheads mark partially or fully unsynapsed autosomes, arrow marks non-homologous SC. Miniaturized images (left bottom corners) show histone H1t, a marker of mid and post-mid pachytene stages (intermediate or high levels, respectively). Roman numbers refer to SC defect types as described in **B**. **E** *Iho1^C7Δ/C7Δ^*cells where either all chromosomes have MLH1 (cell 1) or a synapsed autosome lacks MLH1 (cell 2, arrowhead). **D-E** Bars, 10 µm. See also Figure S5.

Ranking of chromosome axis length suggested that both long and short autosomes were prone to asynapsis in early pachytene (Fig. 5C). In contrast, asynapsis was more frequent for short chromosomes in late pachytene. These observations suggest that, in contrast to short asynaptic chromosomes, long asynaptic chromosomes eventually synapse during pachytene in *Iho1^C7Δ/C7Δ^* spermatocytes. Alternatively, asynapsis of long chromosomes may be more likely to trigger apoptosis, consistent with prior observations that only extensive asynapsis can do so ^53^. Synapsis defects originating from reduced DSB numbers likely underlie both the low testis size (Fig. 3A) and low oocyte numbers (Fig. S3D-E) in *Iho1^C7Δ/C7Δ^* mice, as asynapsis triggers apoptosis in both spermatocytes and oocytes ^52, 54, 55, 64^.

We further tested the formation of crossover-specific recombination intermediates, represented by foci of MLH1, a component of the Holliday junction resolvase ^65–68^. A large fraction of *Iho1^C7Δ/C7Δ^* oocytes (40%, n=68) and spermatocytes (35%, n=100) formed MLH1 foci on all of their chromosomes in pachytene (Fig. 5E, S5F). Nonetheless, defects in MLH1 focus formation were apparent (Fig. 5E-G, S5F-H). *Iho1^C7Δ/C7Δ^* meiocytes always lacked MLH1 foci on chromosomes that were fully unsynapsed (12 autosomes and 25 sex chromosome pairs in spermatocytes, 32 chromosomes in oocytes), and frequently lacked MLH1 foci on partially unsynapsed chromosomes (24 out of 61 chromosomes, quantified only in oocytes, where asynapsis was frequent, Fig. S5D). Interestingly, small fractions of fully synapsed chromosomes of *Iho1^C7Δ/C7Δ^* meiocytes also lacked MLH1 foci in both sexes (3% in oocytes, Fig. S5G, 0.94% autosomes in spermatocytes, Fig. 5F). In spermatocytes, MLH1 focus formation was affected not only on autosomes, but also on sex chromosomes that synapsed their PARs — MLH1 was missing from 28% of PARs in wild type (n=85) and 53% in *Iho1^C7Δ/C7Δ^* (n=74; Fisher exact test, P=0.002), respectively. Accordingly, total MLH1 focus numbers were slightly lower in *Iho1^C7Δ/C7Δ^*than wild-type mice when meiocytes with fully synapsed chromosomes were considered (2 and 1 foci lower medians in males and females, respectively; Fig. 5G and S5H).

One possible interpretation is that the reduced DSB numbers in *Iho1^C7Δ/C7Δ^* meiocytes exceeds the capacity of the as-yet enigmatic crossover homeostasis mechanism, which keeps crossover numbers stable even if numbers of recombination initiation events vary ^69, 70^. Alternatively, the IHO1 C-terminus may support post-DSB functions of IHO1 that promote crossover formation, analogous to functions reported for Mer2 in the fungus *Sordaria macrospora* (*Sm*) ^14^. We favor the former hypothesis because — contrary to *Sm*Mer2, which localizes to SC central regions and CO-specific recombination foci following chromosomal synapsis ^14^ — IHO1 is not detectable on synapsed chromosomal regions ^22, 34^, where CO-generating recombination occurs. Further, conditional depletion of IHO1 soon after most DSBs formed in zygotene does not cause obvious defects in meiosis in mice ^34^. Therefore, post-DSB functions of *Sm*Mer2 are unlikely to be conserved in mammals.

### IHO1_C7Δ does not support accumulation of recombination initiation events in unsynapsed regions

SC formation was suggested to restrict DSB activity to late synapsing genomic regions, where newly formed DSBs are useful for completion of homolog synapsis ^8, 20, 22, 39, 40^. Confirming this hypothesis, delayed synapsis provokes persistence of DSB-machinery ^34, 35, 37^ and accumulation of DSBs ^37–39^ in affected genomic regions in both budding yeast and mice. Asynapsis was frequently observed in *Spo11*-hypomorphic *Tg(Spo11β)^+/-^* spermatocytes, due to an over twofold reduction in total DSB activity as compared to wild type ^39^. Curiously, despite the overall low DSB levels, DSB density reached wild type levels on unsynapsed axes as *Tg(Spo11β)^+/-^*spermatocytes progressed to advanced-late zygotene, where less than 30% of axes were unsynapsed. This phenomenon was attributed to persistent DSB activity in regions where synapsis was delayed.

If, similar to *Tg(Spo11β)^+/-^*, *Iho1^C7Δ/C7Δ^*spermatocytes were able to top up DSBs on asynaptic axes, the relative density of DSB markers on unsynapsed versus synapsed axis (hereafter “unsynapsed-to-synapsed DSB density ratio”) would be the same or higher in *Iho1^C7Δ/C7Δ^*as compared to wild-type in an advanced-late zygotene stage. However, our observations suggested that DSB-factor clusters were not only inefficiently seeded, but were also sensitive to negative-feedback signaling from DSBs in *Iho1^C7Δ/C7Δ^* meiocytes (Fig. 3). Hence, beyond having overall low DSB activity, *Iho1^C7Δ/C7Δ^* meiocytes might fail to efficiently maintain DSB activity on asynaptic axes once DSBs form in leptotene and zygotene.

To test this hypothesis, we examined densities of DSBs on axes by co-staining DMC1 and RPA2, which are preferential markers of recombination foci in unsynapsed and synapsed regions, respectively (Fig. 6A). We compared *Iho1^C7Δ/C7Δ^*with wild type and two DSB-defective genotypes, *Spo11^+/-^* and *Ankrd31^-/-^.* In *Spo11^+/-^* spermatocytes, DSB formation was mildly delayed and reduced (15-30%) relative to wild type ^69^. In *Ankrd31^-/-^* spermatocytes, both DSB-factor cluster formation and recombination initiation were strongly delayed, but DSB focus numbers reached or surpassed wild type levels in late zygotene ^23, 24^. Unsynapsed-to-synapsed DSB density ratios were below one in *Iho1^C7Δ/C7Δ^* spermatocytes (median=0.83) and above 1 in wild-type (median=1.1), *Spo11^+/-^* (median=1.17,) and *Ankrd31^-/-^* (median=1.58) spermatocytes in advanced-late zygotene (Fig. 6B).

**Figure 6.**
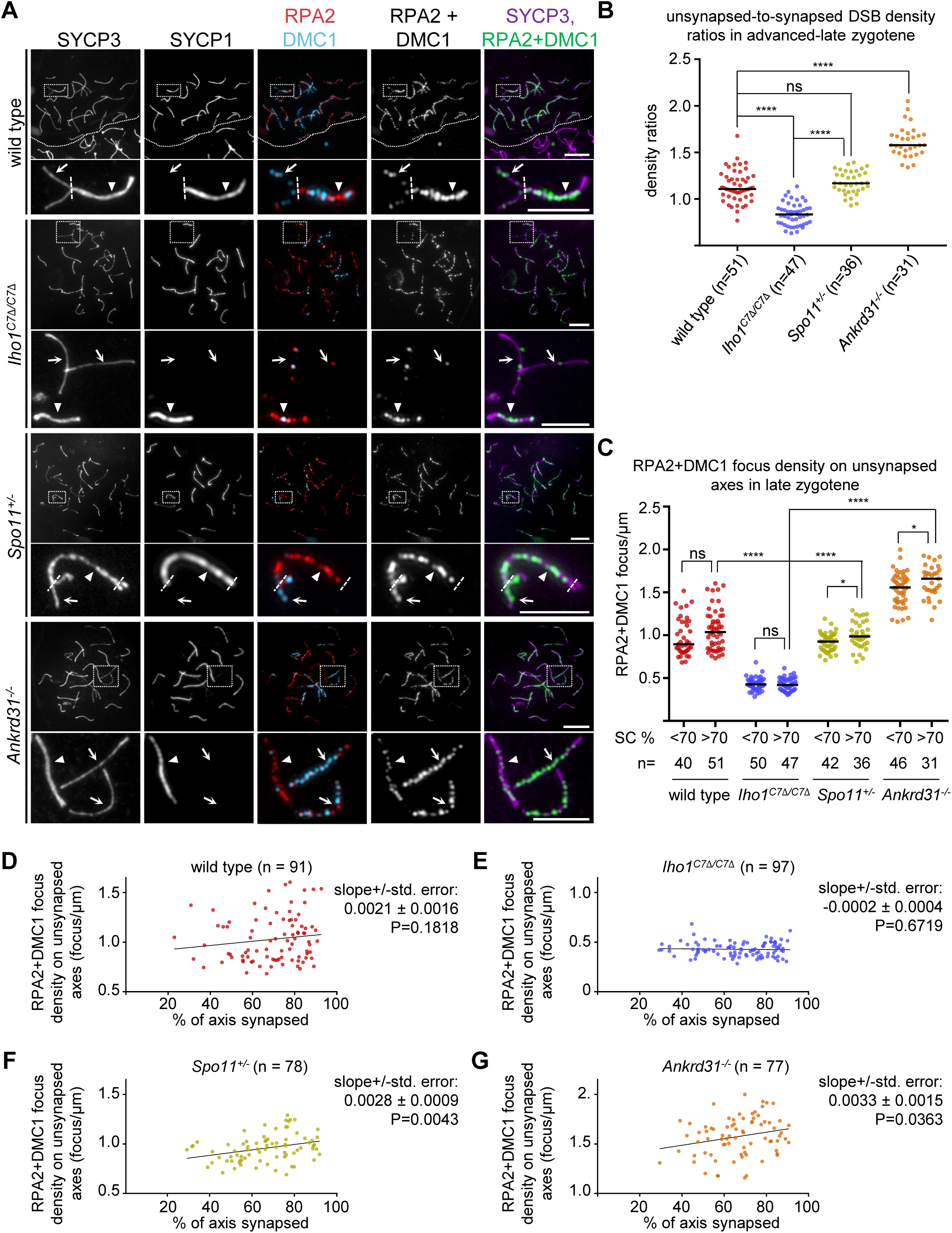
Disruption of IHO1-HORMAD1 interface prevents DSB top-up in late synapsing axes **A** Immunostaining in nuclear spread late zygotene spermatocytes of adult mice. In wild type, dotted line separates a zygotene (top) from a diplotene (bottom) cell. In enlarged insets, dashed lines mark borders between unsynapsed (arrow) and synapsed (triangle) axes. Bars, 10 and 5 µm in low and high magnification panels, respectively. **B** Unsynapsed-to-synapsed DMC1+RPA2 focus density ratios in late zygotene cells where SC formed on >70% of total axis length. **C-G** DSB focus densities on unsynapsed axes in late zygotene cells grouped (**C**, SC formed on <70% or >70% of total axis length) or ordered (**D-G**) according to the extent of synapsis nucleus-wide. **B-G** Each dot represents a single cell. n=numbers of spermatocytes. **B-C** Bars, medians, Mann Whitney U-Test, ns=P>0.05, *=P<0.05, ****=P<0.0001. **D-G** Linear regression (lines), the best-fit slope +/-standard error and the significance of slope deviation from zero (F test, P) are shown.

These observations suggest that late-synapsing regions in *Iho1^C7Δ/C7Δ^*spermatocytes have comparatively low DSB activity. Further, asynaptic regions cannot top up DSBs to levels of early synapsing regions, unlike wild-type, *Spo11^+/-^* and *Ankrd31^-/-^* spermatocytes. Consistent with these conclusions, DSB densities were substantially lower on unsynapsed axes of *Iho1^C7Δ/C7Δ^* than either wild type (∼2.4 fold), *Spo11^+/-^* (∼2.2 fold) or *Ankrd31^-/-^* (∼3.7 fold) spermatocytes in advanced-late zygotene (Fig. 6C-G). Spreading of SC from 20-70% to over 70% of total axis length was not accompanied by a significant increase in DSB densities on unsynapsed axes of late zygotene wild-type or *Iho1^C7Δ/C7Δ^* spermatocytes (Fig. 6C-E). In contrast, a small but significant increase of DSB densities was observed on unsynapsed axes of *Spo11^+/-^*and *Ankrd31^-/-^* spermatocytes as synapsis progressed (Fig. 6C, F-G), which was in line with delayed DSB kinetics in *Spo11^+/-^*and *Ankrd31^-/-^* mice ^23, 24, 69^.

Steadily low DSB densities in *Iho1^C7Δ/C7Δ^* spermatocytes suggest that asynapsis does not enable enduring DSB formation in the absence of IHO1-HORMAD1 interaction. Further, high DSB densities on unsynapsed axes in late zygotene *Ankrd31^-/-^* spermatocytes suggest that, whereas both ANKRD31 and IHO1-HORMAD1 promote DSB-factor clusters on axes, only IHO1-HORMAD1 is required for maintaining DSB activity if synapsis is delayed.

### The IHO1-HORMAD1 interaction and ANKRD31 redundantly enable DSB activity

Not only DSB dynamics (Fig. 6), but also DSB-factor clusters were differentially affected by disruptions of ANKRD31 or IHO1-HORMAD1 (^23–25^ and Fig. 7A-B). Whereas ANKRD31 loss led to disappearance of laDSB-factor clusters in PARs and PAR-like autosomal telomeres ^23, 24^, loss of IHO1 ^24^, HORMAD1 ^25^ or the IHO1 C-terminus (Fig. S3F-G) had no or very little effect on laDSB-factor clusters. Further, initial growth of smDSB-factor clusters was impaired in the absence of ANKRD31, but not in the absence of IHO1 C-terminus or HORMAD1 (Fig. 3E, 3G, 7B).

**Figure 7.**
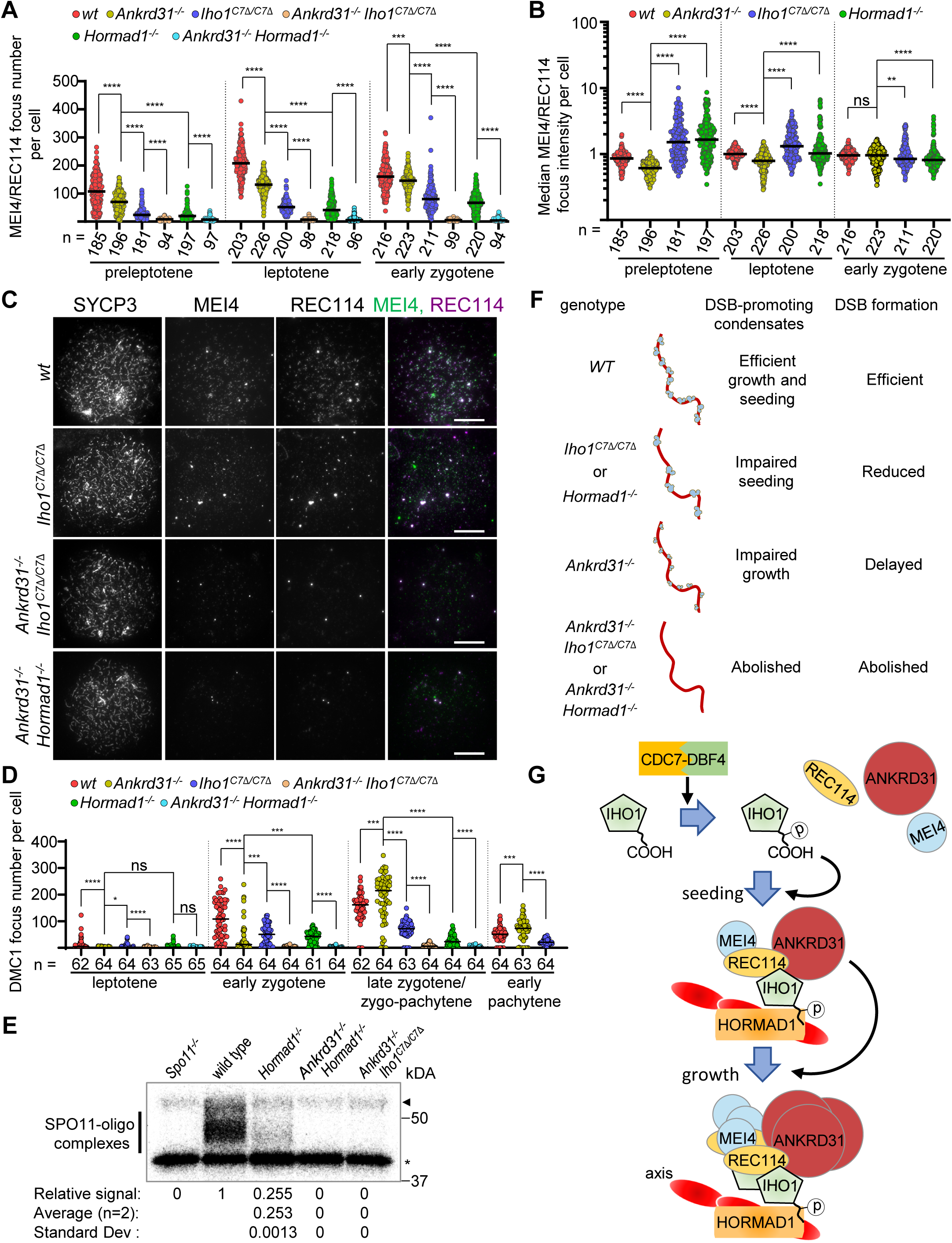
IHO1-HORMAD1 complex and ANKRD31 are redundantly necessary for meiotic DSBs **A-B, D** Numbers of small MEI4-REC114 co-clusters (**A**), MEI4 intensities in MEI4-REC114 co-clusters (**B**, data points show median cluster intensities per cell), and DMC1 focus numbers (**D**) in spermatocytes of adult mice. Zygo-pachytene (**D**, only in SC-defective backgrounds) is equivalent to a mix of late-zygotene and early pachytene stages which are indistinguishable if SC is defective. Bars are medians, n=cell numbers. Mann Whitney U-Test, ns=P>0.05, *=P< 0.05, **=P<0.01, ***=P<0.001, ****=P<0.0001. **C** Immunostaining in nuclear spread leptotene spermatocytes of adult mice. Bars, 10 µm. **E** Radiograph of immunoprecipitated and radioactively labeled SPO11-oligo complexes from testes of adult mice. Bar, SPO11-specific signal, asterisk, nonspecific labelling, and arrowhead, immunoglobulin heavy-chain. Radioactive signals were background-corrected (*Iho1^-/-^* , signal=0) and normalized to wild-type control (1). Means and standard deviations of SPO11-oligo signals are shown from n=2 biological replicates. **F** Schematic summary of phenotypes caused by the disruption of IHO1-HORMAD1 complex and/or ANKRD31. **G** Model for the assembly of DSB-factor clusters on axis. Black arrows represent promotion of (i) IHO1 phosphorylation and promotion of (ii) seeding or (iii) growth of DSB-factor clusters by CDC7-DBF4, IHO1 (in particular, phosphorylated IHO1 C-terminus) and ANKRD31, respectively. See also Figure S6 and S7, Table S6.

ANKRD31 directly interacts with IHO1, REC114 and MEI1 by partially distinct domains (^24^, Fig. S6A, Table S6). Therefore, ANKRD31 may enhance DSB-factor clustering by increasing interconnectivity between cluster components, contrasting a proposed IHO1-HORMAD1 function in anchoring DSB-factor clusters to axes, Accordingly, ANKRD31 and IHO1-HORMAD1 may promote DSB-factor clustering by redundant pathways.

DSB-factor clusters were detected in negligible numbers if both ANKRD31 and either the IHO1 C-terminus or HORMAD1 were disrupted (Fig. 7A, 7C, S6B). DSB-factor clusters resembling PAR-linked clusters of wild type were still present on chromatin of *Iho1^C7Δ/C7Δ^ Ankrd31^-/-^* (92.7%, n=193 cells) and *Hormad1^-/-^ Ankrd31^-/-^* (91.2%, n=136 cells, Fig. 7C, S6B) from preleptotene to early zygotene. However, these clusters rarely associated with PAR-FISH signals (11.4%, n=603 clusters in *Iho1^C7Δ/C7Δ^ Ankrd31^-/-^*, and 12.5%, n=592 clusters in *Hormad1^-/-^ Ankrd31^-/-^*), which may indicate spontaneous aggregation of DSB factors if both IHO1-HORMAD1 and ANKRD31 functions are lost.

In addition, DSB activity was more substantially disabled by coincident disruption of ANKRD31 and the IHO1-HORMAD1 complex. Chromatin-associated levels of ƔH2AX were strongly reduced in *Iho1^C7Δ/C7Δ^ Ankrd31^-/-^* and *Hormad1^-/-^ Ankrd31^-/-^* as compared to *Iho1^C7Δ/C7Δ^*, *Ankrd31^-/-^* and *Hormad1^-/-^* single mutant spermatocytes in leptotene to early zygotene stages (Fig. S6C-D), where ƔH2AX levels principally reflect ATM signaling from DSBs ^71^. *Iho1^C7Δ/C7Δ^ Ankrd31^-/-^* and *Hormad1^-/-^ Ankrd31^-/-^* spermatocytes did not progress beyond a defective zygo-pachytene-like stage where fully formed axes remained largely unsynapsed, yet axes often engaged in multiple apparently nonhomologous entanglements (Fig. S6E). In significant fractions of these spermatocytes, ƔH2AX accumulated in one or a few large chromatin domains (Fig. S6E-F).

These phenotypes closely resembled the phenotypes of *Spo11^-/-^*and *Iho1^-/-^* spermatocytes, where lack of programmed DSBs prevented homology search, and where large ƔH2AX-rich chromatin domains, called pseudo-sex bodies, frequently formed due to ATR signaling from unsynapsed axes and/or sporadic SPO11-independent DSBs ^71, 72^. Consistent with a reported HORMAD1 role in pseudo-sex body formation in DSB deficient meiocytes ^22, 32^, ƔH2AX-rich chromatin domains were less frequent in *Hormad1^-/-^ Ankrd31^-/-^* than *Iho1^C7Δ/C7Δ^ Ankrd31^-/-^*, *Spo11^-/-^* or *Iho1^-/-^* spermatocytes.

Their apparent similarities to *Spo11^-/-^* and *Iho1^-/-^* spermatocytes raised the possibility that *Hormad1^-/-^ Ankrd31^-/-^* and *Iho1^C7Δ/C7Δ^ Ankrd31^-/-^* spermatocytes lacked programmed DSBs. Indeed, DSB foci, as detected by RAD51, DMC1 and RPA2 staining, were rarely, if at all, present in these double mutants (Fig. 7D, S7). Further, SPO11-oligo complexes ^58^ were not detected in testes of *Iho1^C7Δ/C7Δ^ Ankrd31^-/-^* and *Hormad1^-/-^ Ankrd31^-/-^* mice, confirming a lack of DSB activity (Fig. 7E). Therefore, we conclude that ANKRD31 and the IHO1-HORMAD1 complex are redundantly required for meiotic DSB formation and recombination initiation.

## Discussion

Orderly synapsis of homologous chromosomes is enabled by DSB formation on unsynapsed chromosome axes in mammals, yet the mechanism that ensures this appropriate spatial organization of DSB activity is poorly explained. However, recent insights into the mechanism of DNA-bound clustering of DSB factors, together with our data, allows us to propose a model for the axial assembly of the DSB-machinery.

### Enabling seeding and growth of DSB-factor clusters on chromosome axes

Multivalent protein-protein and protein-DNA interactions cooperatively condense the budding yeast orthologs of MEI4 (Mei4), REC114 (Rec114) and IHO1 (Mer2) into DNA-bound co-clusters *in vitro* ^28^. Mer2 also efficiently binds nucleosomes *in vitro* ^29, 73^. DNA binding of DSB factors also shows considerable conservation ^73, 74^. Thus, the *in vitro* clusters of DNA-bound DSB-factors of budding yeast likely model *in vivo* DSB-factor clusters which enable DSB activity in diverse taxa including mammals. However, assembly on DNA cannot fully explain the situation *in vivo*: DSB-factor clusters preferentially assemble on chromosome axes rather than on bulk chromatin ^13, 20, 22, 36, 43, 57^.

Our analysis suggests that ANKRD31 and IHO1-HORMAD1 interaction play crucial and functionally synergistic roles in the axial assembly of the DSB-machinery. Whereas ANKRD31 is not an essential component of DSB-factor clusters ^23, 24^, it directly interacts with at least three essential DSB factors, REC114 ^23, 24^, IHO1 ^24^ and MEI1 (this study). Hence, ANKRD31 may enhance inter-molecular connections between DSB factors during clustering. Accordingly, ANKRD31 supports both the establishment and subsequent growth of DSB-factor clusters.

In contrast, the IHO1-HORMAD1 interaction seems to specifically enable the axial seeding, but not the growth of DSB-factor clusters. The underlying reason could be that IHO1-HORMAD1 interaction axially anchor DSB-factor clusters without influencing their internal architecture. IHO1 recruitment to chromosome axes relies on a stable IHO1-HORMAD1 interaction that does not require other DSB factors. Furthermore, IHO1 requires its conserved C-terminal acidic patch for interaction with HORMAD1, but IHO1 requires neither its C-terminus nor HORMAD1 for interactions with itself, REC114, MEI1 and ANKRD31 (this study) ^22, 24, 75^ indicating distinct molecular requirements for IHO1 interaction with HORMAD1 and DSB-factors. Hence, HORMAD1-mediated assembly of an axial IHO1 platform could enable efficient seeding of cytological DSB-factor clusters by providing arrays of anchor sites for REC114, MEI1 and ANKRD31 recruitment.

There is an ongoing discussion to what extent multivalent low affinity/low specificity interactions versus high specificity interactions contribute to biogenesis of membraneless subcellular compartments ^76^. The paradigm of HORMAD1-IHO1 interaction-promoted seeding of DSB-factor clustering suggests that both high specificity interactions (IHO1-HORMAD1) and multivalent low specificity interactions (DNA-driven condensation) can significantly contribute to mesoscale macromolecular assemblies representing subcellular compartments.

IHO1 could support axial seeding of DSB-factor clusters by distinct mechanisms. It is likely that the dynamics of DSB-factor cluster formation are influenced by a balance between cluster nucleation, dissolution, fusion and growth by incorporation of components from soluble pools whose limited size results in a competition between clusters^28^. Axial IHO1 may directly initiate biochemical nucleation of DSB-factor clusters. A nonexclusive alternative is that DSB-factor clustering is nucleated across chromatin, but that the capture of resultant proto-clusters by axial IHO1 platforms shifts the balance of post nucleation cluster dynamics, thereby stabilizing clusters and/or enabling efficient establishment of cytologically discernible clusters — which likely represent functionally competent DSB machineries. Further, axial IHO1 platforms may attenuate the effect that DSB-triggered negative feedback has on DSB-factor clusters by supporting both existing clusters and/or *de novo* cluster assembly from soluble pools of DSB factors.

The IHO1 C-terminus is not only required for the efficient formation of DSB-factor clusters but also DSBs, indicating that the IHO1-HORMAD1 interaction is a key guarantor of DSBs on unsynapsed axes. Nevertheless, if the axial IHO1 anchor is disrupted by deletion of IHO1 C-terminus or HORMAD1, functional DSB-factor clusters and dependent DSBs still form on axes, albeit at low frequency. One or more DSB factors and axis components may engage in cooperative low affinity interactions, which permits seeding of cytological DSB-factor clusters with low efficiency. Due to their low numbers, successfully seeded clusters may grow bigger than normal by sequestering soluble DSB factors. These DSB-factor clusters are strictly dependent on ANKRD31, which may reflect a need for high valency of interactions between DSB factors to allow stabilization of their heteromeric clusters in the absence of an axial IHO1 platform.

LaDSB-factor clusters on PARs and PAR-like regions do not require HORMAD1 or IHO1, but critically depend on ANKRD31, REC114 and MEI4 ^21, 23–25^. Thus, instead of a stable IHO1-HORMAD1 interaction, cooperative protein-protein and protein-DNA interactions may take the lead in seeding laDSB-factor clusters, resembling *in vitro* condensate formation. PARs and PAR-like regions are rich in mo-2 minisatellite repeats ^25^, which may provide arrays of binding sites for efficient seeding of DSB-factor clusters independent of IHO1-HORMAD1 interaction.

### IHO1-HORMAD1 interaction supports error correction of synapsis

Delays in homolog synapsis — which occur in several mouse models, i.e. *Ankrd31^-/-^*, *Tg(Spo11β)^+/-^* ^39^ or *Spo11^+/-^* — are thought to be remedied by persistent DSB activity on asynaptic axes ^37–40^. Curiously, *Iho1^C7Δ/C7Δ^* meiocytes show enduring asynapsis without associated accumulation of DSBs, indicating that maintenance of DSB activity in asynaptic regions depends on IHO1-HORMAD1 interaction. The IHO1-HORMAD1 interaction seems to enhance not only the formation of DSB-factor clusters, but also their resistance to destabilizing negative feedback from previously formed DSBs after leptotene. We hypothesize that loss of the latter function considerably contributes to synapsis defects in *Iho1^C7Δ/C7Δ^*meiocytes by preventing asynapsis-enabled maintenance of DSB activity and downstream correction of synapsis errors.

### Evolutionary comparison suggests that axial localization of DSB-machinery enables coordination of DSB activity and homolog pairing

There is considerable evolutionary divergence in the sequences and functions of DSB factors and the spatiotemporal control of DSBs. Co-evolution of distinct features of DSB-machinery and DSB regulation may indicate their functional coupling, hence inter-taxa comparisons reveal general principles of DSB control.

Interactions between HORMA domain proteins and Mer2/IHO1-family proteins are thought to provide axial anchors for the DSB-machinery in multiple species ^20, 22, 29–31, 36, 43^. Yet, redundancies in mechanisms that link DSB-machinery to axes diminish the importance of the interaction between Mer2/IHO1- and HORMAD1-family proteins in some taxa ^30, 36^. Consistent with this principle, a HORMAD1 ortholog has not been reported in the fungus *Sordaria macrospora*, and an acidic patch is absent from the C-terminus of the *Sm*Mer2 ^14^. Furthermore, *Sm*Mer2 exhibits unique behavior among Mer2/IHO1-family proteins by being present on chromosomes from early meiosis till early post-meiotic divisions and by having post-DSB roles in homolog pairing, recombination and late meiotic chromosome compaction in addition to being essential for DSBs. In contrast, Mer2/IHO1-family proteins of all other examined taxa diminish from chromosomes following synapsis and likely lack major functions besides DSB formation ^18, 22, 31, 34, 36, 43^. Thus, while *Sm*Mer2 retained its axis–associated role in enabling DSB formation, it seemingly acquired new post-DSB roles that co-evolved with (i) an altered control of its chromatin binding, (ii) the loss of its C-terminal acidic patch, and (iii) a presumed lack of a HORMAD1 ortholog.

A unique paradigm is presented by the nematode *Caenorhabditis elegans* (worm hereafter), which lacks a recognizable Mer2/IHO1 ortholog but possesses REC114 (DSB-1 and 2) and MEI4 (DSB-3) orthologs ^14, 15^, which are crucial for DSBs ^15, 77, 78^. Unlike DSB factors of most other taxa, worm DSB-1/2/3 form clusters on bulk chromatin instead of axes ^15, 77, 78^. The absence of a Mer2/IHO1-family protein may deprive DSB-factor clusters from an axial anchor, thereby preventing their axial enrichment. Judging from the localization of DSB-factors, it is conceivable that DSBs form off-axis in worms ^15^. Yet, recombination foci seemingly associate with the axis ^79^, suggestive of a post-DSB mechanism that recruits recombination intermediates to axes ^80^. Consistently, irradiation-induced DSBs — which are not confined to axes — give rise to axis-associated recombination foci in mice, and a similar configuration may be present in irradiated worms ^34, 63, 81^. The existence of post-DSB mechanisms for axial recruitment of recombination foci raises the question why DSB activity is focused on axes in most taxa.

In most taxa, recombination initiation by DSBs is required for synapsis of homolog axes. Therefore, generation of DSBs — which are potentially genotoxic — is not beneficial once synapsis is achieved. Axial association places DSB-factors in the physical context of the SC, allowing DSB activity to be controlled by synapsis. DSB formation is maintained in unsynapsed regions, where DSBs are needed for promoting synapsis, whereas synapsis depletes DSB-factor clusters and ends DSB activity ^34, 35, 37–39^. Unlike most studied taxa, worms synapse homolog chromosomes using pairing centers rather than DSBs ^82–84^. Nevertheless, worms still require DSBs for CO formation ^84^. Due to altered requirements, worms maintain competence for DSB formation on synapsed chromosomes, and they link depletion of DSB-factor clusters and cessation of DSB activity to the formation of COs instead of synapsis ^77, 78^.

We speculate that axial accumulation of DSB factors is not conserved in worms because unique alterations in homolog pairing mechanisms make it unnecessary to coordinate DSB activity with synapsis in this taxon. Accordingly, one reason why organisms that rely on DSBs for synapsis may accumulate DSB-factors on the axes — besides priming axial localization of recombination intermediates — is that axis association of DSBs enables appropriate coordination of DSB activity with homolog synapsis, which both aids efficient homolog pairing and prevents excessive DSBs during meiosis.

## ACKNOWLEDGMENTS

We thank Dunja Knapp for insightful comments and proofreading the manuscript, M. Munzig for lab support, C. Santocanale for advice with DDK inhibitors and DDK assays, R. Jessberger for departmental support and sharing ideas and reagents (anti-SYCP3 antibody), E.L. Huttlin and S.P. Gygi for sharing mass spectrometry data and information about IHO1 phosphorylation, B. De-Massy, M. Biot and C. Grey for sharing ideas and unpublished data about genomic localization of DSB factors. I. D., V. T., F. P., K. R., M.S., N.S.I. and A. T. were supported by the Deutsche Forschungsgemeinschaft (DFG; grants: TO421/5-1, TO421/6-1/2, TO421/7-1, TO421/8-1/2, TO421/10-1, TO421/11-1, TO421/12-1 and TO421/14-1), HFSP research grant RGP0008/2015 and core funding from the Faculty of Medicine at the TU Dresden. Work in the S.K. laboratory was supported in part by US National Cancer Institute Cancer Center support grant P30 CA08748 and US National Institutes of Health grant R35 GM118092. J.R.W. and E.S. were funded by the Max Planck Society, and the DFG grant number WE 6513/2-1. B.N. and F.H. were funded by the GFF-NÖ (Stiftungsprofessur). We also acknowledge Grant “Gruppi di Ricerca 2020” from Regione Lazio, Italy (n. A0375-2020-36618) to MB, and Fondo di Beneficienza Intesa Sanpaolo (n. B/2021/0228) to MB. The Core Facility was generously supported by grants from European Regional Development Fund (ERDF/EFRE) (Contract #100232736) and the Deutsche Forschungsgemeinschaft (DFG) grants (INST 269/731-1 FUGG).

## AUTHOR CONTRIBUTIONS

I.D.: conceptualization, methodology, investigation, formal analysis, visualization, writing– original draft preparation, data curation and project administration.

V.T., F.P., K.R., J.X., M.Bo., B.N., N.S.I., E.S., H.T.E., S.Y., T.G., M.G., A.B.: investigation, formal analysis.

M.Sta.: conceptualization, investigation, formal analysis. M.Ste.: supervision, writing–review and editing.

M.Ba., A.S., J.R.W., F.H., S.K.: supervision, writing–review and editing, funding acquisition.

A.T.: conceptualization, writing–original draft preparation, review and editing, supervision, project administration, funding acquisition.

## EXPERIMENTAL MODEL AND SUBJECT DETAILS

### Animal experiments

Gonads were collected from mice after cervical dislocation. Most cytological experiments of spermatocytes were carried out on samples collected from adult mice unless indicated otherwise. We used testes of juvenile mice (12-13 days old) for most biochemical experiments to enrich pre-pachytene spermatocytes and to ensure that the cellular compositions of testes were similar in wild-type and meiotic recombination mutant mice. The mid pachytene stage, where most recombination defects trigger apoptosis ^52^, is reached by a majority of spermatocytes at around 14 days of age during the first developmental wave of meiosis in mice. Therefore, testes are directly comparable in wild-type and meiotic mutant mice at 12-13 days of age.

Animals were used and maintained in accordance with the German Animal Welfare legislation (‘‘Tierschutzgesetz’’). All procedures pertaining to animal experiments were approved by the Governmental IACUC (‘‘Landesdirektion Sachsen’’) and overseen by the animal ethics committee of the Technische Universität Dresden. The license numbers concerned with the present experiments with mice are TVV 2014/17, TV A 8/2017, TV A 23/2017, and TV vG 3/2022.

### Generation of *Iho1^C7Δ^* mice

*Iho1^C7Δ^* mutant line was generated using CRIPSR/Cas9 genome editing ^85^, targeting exon7 of *Iho1* gene. A mixture of gRNA: GGATTTTGATAGCAGCGATGATA (12.5 ng/ml, IDT), designed using the online platform at http://crispr.mit.edu/guides and https://gt-scan.csiro.au/gt-scan ^86^, and Cas9 nuclease mRNA (50 ng/ml), prepared as described before ^24^, was injected into pronucleus/cytoplasm of fertilized oocytes. The oocytes were subsequently transferred into pseudopregnant recipients. A founder mouse that was heterozygote for an insertion of T nucleoside that caused a premature stop codon was bred with C57BL/6JCrl wild-type mice to establish mouse lines. All experiments reported in the manuscript are based on samples from mice that were derivative of this founder mouse after at least three backcrosses.

## METHOD DETAILS

To prepare Cas9 mRNA, we first used the restriction enzyme PmeI to linearize the plasmid MLM3613 (Addgene #42251) ^85^ that harbors a codon optimized Cas9 coding sequence and a T7 promoter for Cas9 mRNA *in vitro* synthesis. We then used the linearized MLM3613 as template to synthesize the 5′ capped and 3′ polyA-tailed Cas9 mRNA using the mMESSAGE mMACHINE® T7 Ultra Kit (ThermoFisher, cat no: AM1345) according to the manufacturer’s instructions.

### Genotyping

Tail biopsies were used to generate genomic DNA by overnight protease K digestion at 55°C in lysis buffer (200mM NaCl, 100mM Tris-HCl pH 8, 5mM EDTA, 0.1% SDS), followed by heat inactivation for 10 min at 95C. Genotypes were identified either by heteroduplex mobility assay (HMA) or by the use of mismatch primers in genotyping PCRs. For HMA, PCRs were carried out with CATGACCACCAGAAGCGTCA and AATGTTTTCACCAAGGACATAC primers, which annealed to sequences flanking the mutated site in the Iho1 gene. Genotyping PCR products or one-to-one mixtures of genotyping PCRs and PCRs of wild type genomes were supplemented with EDTA in 10mM final concentration. After a 2-minute denaturation cycle at 98 °C and a 30-minute annealing cycle at 25°C, electrophoresis was performed in 8% polyacrylamide gels that were prepared with TBE buffer. As an alternative strategy to HMA, and to allow analysis of genotyping PCRs on agarose gels we also carried out PCRs with primers that annealed to the site of mutation in the *Iho1* gene. Due to only a single base difference in the sequence of wild-type and mutant loci, primers that annealed to the site where the mutation was introduced enabled amplification from both wild-type and mutant loci. To better distinguish between wild-type and mutant templates we introduced mismatches into both wild-type and mutant specific primers. CAGAGATCAAAGAGAGGTGG and CATCGCTGCTATCAA<u>G</u>ATC (mismatch nucleotide underlined) primers allowed high specificity amplification of a 385 bp product from the wild type allele. CAGAGATCAAAGAGAGGTGG and CATCGCTGCTATCAAAA<u>G</u>TC (mismatch nucleotide underlined) allowed high specificity amplification of a 386 bp product from the *Iho1^C7Δ^* allele.

### Testis electroporation

To overexpress tagged wild type and mutant proteins in spermatocytes, we injected 6–8 μl of an expression vector (3-5 μg/μl) under the control of a CMV promoter into the rete testis of live juvenile mice (13 days postpartum/dpp) according to published protocol ^87, 88^. 45 minutes after injection, testes were held between tweezer type of electrodes (CUY650P5, Nepagene) and *in vivo* electroporation was carried out with four 50-ms pulses (35 Volts) with 950 ms intervals, then four equivalent pulses with opposite polarity (NEPA21 Electroporator, Nepagene).

### Protein extracts, immunoprecipitation and western blotting

#### Total protein extraction

Testes of 8dpp ─ in the case of testis cell cultures ─ or 12-13 dpp juvenile mice were collected, and tunica albuginea was removed. Extracts were prepared from testes either after *in vitro* culture (two days of culture with or without CDC7 inhibitors, see details later), or testes were immediately used, or they were flash frozen in liquid nitrogen for later use. Frozen testes were thawed on ice for 10-15 minutes before use. Fresh or thawed tissue were homogenized with the help of disposable tissue grinder pestle (VWR, BELAF199230000) in resuspension buffer (50 mM Tris pH=7.4 150 mM NaCl, supplemented with protease and phosphatase inhibitors: 1 mM Phenylmethylsulfonyl Fluoride (PMSF), complete EDTA-free Protease Inhibitor Cocktail tablets (Roche, 11873580001), 0.5 mM Sodium orthovanadate, Phosphatase inhibitor cocktail 1 (Sigma, P2850) and Phosphatase inhibitor cocktail 2 (Sigma, P5726), protease and phosphatase inhibitor cocktails were used at concentrations recommended by the manufacturers). Testes homogenates were mixed 1:1 ratio with 2X lysis buffer (Tris 50 mM pH=7.4, 850 mM NaCl, 2% Triton X-100, 2% NP40, 1% Sodium deoxycholate/NaDOC, 20 mM MgCl2, supplemented with protease and phosphatase inhibitors as above). Testis homogenates were lysed with the help of overhead rotator for 60 min at 4C in the presence of benzonase (Merck Millipore) to digest DNA during lysis. Total testis lysates were mixed with 2x Laemmli sample buffer (with 10% b-Mercaptoethanol) 1:1 ratio and incubated for 10 min at 95C. Samples were run on 10% gel with using 10 × 10.5cm gel cassette (Cytiva miniVE Vertical Electrophoresis System) to improve separation of phosphorylated and non-phosphorylated IHO1 bands.

#### Fractionation of testis extracts based on Triton X-100 solubility

Testes were homogenized with the help of disposable tissue grinder pestle in resuspension buffer (as described for total extracts). Big tissue chunks were additionally disrupted by 200 µL pipette tip with a cut end. Homogenized testes were divided into 2 parts – ¼ of the homogenate for total extract and ¾ for fractionation. Both aliquots were mixed 1:1 with 2x Triton X-100 buffer (Tris 50 mM pH=7.4, 150 mM NaCl, 0.6 % Triton X-100, supplemented with protease and phosphatase inhibitors) and incubated at 4°C for 30 min, constantly mixed with overhead rotator. Thereafter, lysed homogenates intended for total extracts were placed on ice until further processing, and aliquots intended for fractionation were centrifuged at 4°C for 10 min at 16000g. Supernatants were collected to a new tube as soluble fraction. Pellets containing insoluble fractions were resuspended in the same buffer in the same volume as the supernatant. The unfractionated extracts and both the Triton X-100 soluble and insoluble fractions were mixed with 2x Lysis buffer (850mM NaCl, 50mM Tris-HCl pH 7.5, 2% Triton-X100, 2% NP-40, 1% NaDOC, 1mM MgCl2 supplemented with protease inhibitors and benzonase). After 1h of incubation, unfractionated extracts and soluble and insoluble fractions were mixed with 2x Laemmli sample buffer (with 10% b-Mercaptoethanol) 1:1 ratio and incubated for 7 min at 95°C.

#### IHO1 dephosphorylation assay

Total testis lysates were prepared as described above but phosphatase inhibitors were omitted. Each pair of testes from one juvenile male mouse was lysed in 50 µL 1x lysis buffer. After 1.5h of incubation sample was centrifuged 16 000g for 10 min. Then supernatant was diluted 2 times with phosphatase dilution buffer: 100mM NaCl, 50mM Tris-HCl pH 7.5 supplemented with protease inhibitors. Afterwards, lysate was split into 4 aliquots (35 µL each). All aliquots were supplemented with 10X NEBuffer for Protein MetalloPhosphatases (PMP) and 10X MnCl2 (NEB). The first aliquot (Untreated control) was supplemented with phosphatase inhibitors (PPase inhibitors): 0.5 mM sodium orthovanadate, phosphatase inhibitor cocktail 1 (Sigma, P2850) and phosphatase inhibitor cocktail 2 (Sigma, P5726). They were used at concentrations recommended by the manufacturers. After that 2x Laemmli buffer (with 10% b-Mercaptoethanol) was added and sample was boiled 95°C for 7 min. The second (PPase inhibitors only) and third (PPase inhibitors + Lambda PPase) aliquots were also supplemented with PPase inhibitors. 2 µL of Lambda Protein Phosphatase (NEB, P0753S) was added to aliquots 3 and 4 (PPase only). Final volume of each sample was 50 µL. Aliquots 2-4 were incubated at 30°C for 1.5 h. After incubation, 2xLaemmli buffer (with 10% b-Mercaptoethanol) was added 1:1 ratio and samples were incubated at 95°C for 7 min.

#### IHO1 phos-tag gel preparation and Western Blot

For analysis on phos-tag gel, total lysates of testis were prepared as described above. For preparation of phos-tag gels the following reagents were used (indicated final concentrations): 8% acrylamide/bis-solution (ROTH), 20µM phos-tag acrylamide AAL-107 aqueous solution (Fujifilm), 40mM MnCl2 (Sigma), 0.04% SDS (ROTH), 0.04% APS and TEMED (Bio-Rad)(ROTH), 375 mM Tris-HCl pH 8.8 (ROTH). Following electrophoresis, gels were incubated 3x 10 minutes in Tris/Glycine transfer buffer without methanol and supplemented with 10mM EDTA. After treatment, proteins were transferred on PVDF-membrane with Tris/Glycine transfer buffer supplemented with 20% Methanol for 2.5h (300mA). PVDF-membranes were blocked for 1h with 5% non-fat dry milk in TBS-T, followed by incubation overnight with primary antibodies.

#### IHO1 and HORMAD1 immunoprecipitation

Total testis lysates were spun down for 10 min at 16.000g after lysis. Supernatants were collected to a new tube and diluted two times with 1x IP lysis buffer (50 mM Tris-HCl pH 7.4, 500 mM NaCl, 1% Triton X-100, 1% NP40, 0.5% NaDOC, supplemented with protease and phosphatase inhibitors). To reduce non-specific immunoprecipitation we precleared lysates by adding 1mg Dynabeads Protein A (Invitrogen), and mixing them for 4 hours at 4 °C in an overhead rotator. After pre-clearing, protein extracts were incubated with affinity beads at 4°C overnight. Either rabbit anti-IHO1 (3μg) antibodies or rabbit anti-HORMAD1 (3μg) antibodies were cross-linked to 1.5mg of Dynabeads Protein A (Invitrogen) using 20 mM Dimethyl Suberimidate according to standard protocols to prepare affinity beads. Following overnight incubation in the protein extracts, affinity beads were washed three times with 1x IP lysis buffer. Immunoprecipitated materials were eluted by incubating the beads in 50ul 1x Laemmli sample (supplemented with 5% b-Mercaptoethanol) buffer for 10min at room temperature, agitated 2-3 times with vortex in between. The resulting eluates were analyzed with SDS-PAGE and immunoblotting using standard methods. Briefly, proteins from extracts were separated on SDS polyacrylamide gels and blotted onto Nitrocellulose membrane (Hybond ECL Cat#RPN2020D, GE Healthcare). Membranes were blocked using Skimmed Milk 5%, 0.05% Tween, TBS blocking solution. Band intensities were measured using ImageJ.

##### Expression and purification

Codon-optimized synthetic dsDNA (Geneblocks) for IHO1 and HORMAD1 were synthesized by IDT (Leuven, Belgium). Constructs were cloned into pCOLI or pLIB vectors for E.coli or insect cell expression, respectively ^89, 90^.

For insect cell expression, virus was generated from Sf9 cells using standard protocols. All the IHO1 truncations were expressed with the 3C HRV cleavable N-terminal MBP tag in suspension culture of High Five™ Cells (Invitrogen) in Sf-900™ III SFM medium, infected with virus (diluted 1:000), for 48 h at 27°C. Cell pellets were washed with 1× PBS and resuspended in lysis buffer (50 mM HEPES pH 7.5, 300 mM NaCl, 5% glycerol, 0.1% Triton-X 100, 1 mM MgCl_2_, 5 mM β-mercaptoethanol, AEBSF 25 μg/mL). For cell lysis, sonication at 30% of power was used, and lysate was cleared by centrifugation at 40000 g for 40 min. Cleared lysate was incubated with benzonase (Sigma-Aldrich) at 8°C for 20 min and then applied on a 5 ml MBP-trap column (Cytiva), wash with lysis buffer and eluted with 1 mM maltose solution based on lysis buffer. The elution fractions with highest content of IHO1 truncations were applied to Resource Q column (GE healthcare) equilibrated with (50 mM HEPES pH 7.5, 100 mM NaCl, 5% glycerol, 1 mM TCEP). The proteins were eluted with the gradient of NaCl up to 1 M. Proteins after ion-exchange chromatography were concentrated on the Pierce^TM^ Protein Concentrator PES (30K MWCO, 5-20 ml) and applied to the Superdex200 10/300 Increase column (Cytiva) equilibrated with 50 mM HEPES pH 7.5, 300 mM NaCl, 5% glycerol, 1 mM TCEP.

SUMO-His N-terminally tagged HORMAD1 HORMA (amino acids 1-235) was expressed in chemically competent E.coli BL21(DE3) cells overnight at 18°C with induction of protein expression by 250 µM IPTG. The subsequent purification of the protein was carried out according to the same protocol as with IHO1 truncations, except for the affinity chromatography step. For this cleared lysate of the cells was applied to the Talon column (Cytiva), equilibrated with the lysis buffer (50 mM HEPES pH 7.5, 300 mM NaCl, 10% glycerol, 1 mM MgCl_2_, 0.1% Tween20, 5 mM imidazole, 5 mM β-mercaptoethanol), wash with two steps of lysis buffer with higher concentrations of imidazole (10 and 20 mM) and eluted with lysis buffer containing 250 mM imidazole.

##### Amylose pulldown assay

Amylose pulldown assay was performed with Amylose Sepharose beads (New England BioLabs) in a pulldown buffer (20 mM HEPES pH 7.5, 100 mM NaCl, 5% glycerol, 1 mM TCEP, 0.1% Tween20). The beads were pre-blocked with 1 mg/ml BSA in pulldown buffer for 2 h at 8°C washed twice with 500 µL and then resuspended in equal volume of pulldown buffer. The MBP-tagged IHO1 truncations (residues 359-574, 440-574, 358-567, 439-56) were incubated at 1 µM concentration as baits with 12 µM of SUMO-His tagged HORMAD1 HORMA (residues 1-235) for 30 min on ice. Input samples were mixed with 6x SDS-PAGE sample buffer and loaded on the 10% SDS-PAAG (5 μL for anti-MBP and anti-HORMAD immunostaining and 10 μL for Coomassie staining). Resuspended beads were incubated with protein mix on shaker at 8°C for 30 min and then washed twice with 100 μL of the pulldown buffer. Elution was performed by adding 40 μL of 1x SDS-PAGE sample buffer (10 μL was loaded for each immunostaining and 20 μL for Coomassie staining). After SDS-PAGE, one gel was stained with Coomassie Blue (Blaue Jonas Stain, GRP GmbH) and proteins from another gel were transferred onto nitrocellulose membrane. After Western blotting the membrane was blocked in 5% nonfat milk in PBS with 0.1% Tween-20 (PBST) and incubated with primary antibodies overnight at 4°C. Rabbit polyclonal anti-HORMAD1 antibodies (Abcam) were used at the concentration 1 μg/μl. After incubation with primary antibodies, the membrane was washed 3 times with excessive amount of PBST, incubated with secondary anti-rabbit IgG antibodies and washed with PBST. Detection of the proteins was performed using ECL Prime Western Blotting Detection Reagent.

### Analysis of DDK-mediated phosphorylation of IHO1 C-terminal peptide

#### *In Vitro* phosphorylation of IHO1 peptides by DDK

C-terminal peptides of IHO1 (NLLCDPDFDSSDDNF ending with either NH2 or COOH groups) were added in 0.4mM concentration to 100-μl reaction mixes containing 1x reaction buffer, 100 mM DTT, 10mM Ultrapure ATP and 0.7 μg of recombinant full-length human CDC7/DBF4 kinase (Promega, V5088). Reactions were incubated at 37 °C for 120 min and then terminated by freezing on dry ice before mass spectrometry.

#### Phospho peptide preparation and SEC separation for mass spectrometry

Frozen samples were thawed and 29 µL ACN with 0.43% TFA was added to 100 µL of sample to achieve a concentration of 22.5% ACN and 0.1% TFA in total. The samples were left on room temperature for 5 min and centrifuged at 14 000 g for 10 min to remove possible precipitation products. 16 µL of the sample was in injected into a Dionex Ultimate 3000 (Thermo Fisher Scientific, Vienna, Austria) using a Cytiva Superdex 30 Increase Precision 3.2/300 GL Prepacked SEC Column (Cytiva, Marlborough, MA, USA), 77.5% H_2_O, 22.5%, 0.1% TFA as mobile phase and was separated with a flow rate of 25 µL min^-1^. Absorption of UV light at 215 nm detected a major peak at 56.9 min corresponding to a molecular mass of about 1500-2000 Da. The collected fraction at 54-60 min was dried and used for HPLC-MS analysis.

#### HPLC-MS analysis

For the identification of the peptides a Dionex Ultimate 3000 RSLCnano System coupled to an Orbitrap Eclipse Tribrid Mass Spectrometer (both Thermo Fisher Scientific, Vienna, Austria) was used.

The dried samples were dissolved in 25 µL mobile phase A (98% H_2_O, 2% ACN, 0.1% FA), shaken for 10 min at 30 °C and centrifuged at 14000 g for 1 min. 2 µL were injected onto a PepMap RSLC EASY-Spray column (C18, 2 µm, 100 Å, 75 µm x 15 cm, Thermo Fisher Scientific). Separation occurred at 300 nL min^-1^ with a flow gradient from 2-35% mobile Phase B (2% H_2_O, 98% ACN, 0.1% FA) within 25 min resulting in a total method time of 55 min. Mass spectrometer was operated with the FAIMS Pro System in positive ionization mode at alternating CV -60 and -75. The scan range was 350-2000 m z^-1^ using a resolution of 60,000 @200 m z^-1^ on MS1 level. Isolated peptides were fragmented using CID at a collision energy of 30% and fragments were analyzed in the Orbitrap with a resolution of 30,000 @200 m z^-1^.

#### Peptide identification

Peptides were identified using Proteome Discoverer 2.5 (Thermo Fisher Scientific) employing the Sequest HT search engine ^91^. Phosphorylation sites were determined using the implemented ptmRS node. Raw files were searched against the mouse proteome derived from UniProt (*Mus musculus*, 17082 reviewed entries, canonical and isoforms) and the additional C-terminal Iho1 peptide NLLCDPDFDSSDDNF. Phosphorylation (STY) and Amidation (C-terminal) were set as variable modifications.

#### Spo11-oligo

To measure levels of SPO11-oligonucleotide complexes in testes, we carried out SPO11 immunoprecipitations and SPO11-oligonucleotide detection as published previously ^32, 58^. Both testes from one adult mouse were used for each experiment.

#### Yeast two-hybrid (Y2H) assay

Yeast two-hybrid experiments with mouse proteins were performed as described previously with minor modifications ^22^. Pairwise interactions were carried out in the Y2HGold Yeast strain (Cat. no. 630498, Clontech). For transformation of Y2HGold with bait and prey vectors, yeast were grown in 2xYPDA medium overnight at 30°C, 200 rpm shaking. Afterward, yeast cells were diluted to 0.4 optical density (OD, measured at 600 nm) and incubated in 2x YPDA for 5 h at 30C, 200 rpm shaking. Cells were harvested, washed with water and re-suspended in 2 mL of 100 mM lithium acetate (LiAc). 50 µL of this cell suspension was used for each transformation. Transformation mix included 1 µg of each vector (bait and prey), 60 µL of polyethylene glycol 50% (w/v in water), 9 µL of 1.0M LiAc, 12.5 µL of boiled single-strand DNA from salmon sperm (AM9680, Ambion), and water up to 90 µL in total. The transformation mix was incubated at 30°C for 30 min, and then at 42C for 30 min for the heat shock. The transformation mix was removed following centrifugation at 1000 g for 10 min, and then cells were resuspended in water, and plated first on -Leu -Trp plates to allow selective growth of transformants. After 2-3 days of growth, transformants were plated both on -Leu -Trp and -Leu -Trp -Ade -His plates for 2-7 days to test for interactions. We followed the manufacturer’s instructions for media and plate preparation. Yeast two-hybrid essays of Arabidopsis proteins were performed as described previously ^92^.

### Immunofluorescence microscopy

#### Preparation of spermatocyte spreads

Preparation of nuclear surface spreads of spermatocytes was carried out according to earlier described protocols with minor modifications ^24, 93^. Testes were minced with scalpels in PBS pH 7.4 and collected into a clean tube. The testis suspension was left standing for few minutes to allow sedimentation of large seminiferous tubule fragments. Supernatant was collected mixed with hypotonic extraction buffer (30mM Tris HCl pH 8.2, 50mM sucrose pH 8.2, 17mM sodium citrate pH 8.2, 5mM EDTA pH 8.2, 0.5mM DTT) in 1:1 ratio and incubated for 5 minutes at room temperature. Cell suspensions were diluted three times in PBS pH 7.4 and were centrifuged for 5 min at 1000 g. Supernatants were discarded and pelleted cells were resuspended in PBS pH 7.4. Cell suspensions were mixed 1:2 with 100mM sucrose solution immediately prior to fixation. Cell suspensions were added to three to five times higher volume droplets of filtered (0.2 mm) 1% paraformaldehyde (PFA), 0.15% Triton X-100, 1mM sodium borate pH 9.2 solution on diagnostic slides, and incubated for 70 min at room temperature in wet chambers. Then, slides were air dried either on the bench or in a fume hood. Finally, the slides were rinsed by water, incubated in 0.4% Photo-Flo 200 (Kodak) for 5-10 min, rinsed 3 times with distilled water, and dried at room temperature before proceeding to immunostaining.

#### Preparation of oocyte spreads

To prepare nuclear surface spread oocytes, two ovaries from each mouse were incubated in 20 µL hypotonic extraction buffer at room temperature for 15 min (Hypotonic Extraction Buffer/HEB: 30 mM Tris-HCl, 17 mM Trisodium citrate dihydrate, 5 mM EDTA, 50 mM sucrose, 0.5 mM DTT, 0.5 mM PMSF, 1x Protease Inhibitor Cocktail). After incubation, HEB solution was removed and 16 µL of 100 mM sucrose in 5mM sodium borate buffer pH 8.5 was added. Ovaries were punctured by hypodermic needles (27Gx1/2“) to release oocytes. Big pieces of tissue were removed. 9 µL of 65 mM sucrose in 5 mM sodium borate buffer pH 8.5 was added to the cell suspension and incubated for 3 min. After mixing, 1.5 µL of the cell suspension was added in a well containing 20 ml of fixative (1% paraformaldehyde, 1mM borate buffer pH: 9.2, 0.15% Triton X-100) on a glass slide. Cells were fixed for 45 min in humid chambers, and then slides were air-dried. Upon completion of drying slides were washed with 0.4% Photo-Flo 200 solution (Kodak, MFR # 1464510) for 5 min and afterward they were rinsed with distilled water temperature before proceeding to immunostaining.

##### Staining procedures of nuclear spreads

Previously described blocking and immunostaining procedures ^22^ were optimized for immunostaining with each combination of antibodies; details are available upon request. In most cases, slides were treated with blocking buffer (2% BSA, 0.02% Na azide, 0.05% Tween20 and 0.05%Triton X-100, PBS, pH7.4 plus 0.05% Na azide) for 30 minutes before incubation with primary and secondary antibodies in blocking buffer. Combined immunostaining and PAR-FISH was performed on spermatocyte spreads as before ^24^.

#### Immunofluorescence on gonad sections

Testes were collected with care to ensure minimal disturbance of internal structure, and immediately fixed for 40 minutes at room temperature in 3.7% formaldehyde, 100mM sodium phosphate pH 7.4, 0.1% Triton-X. After fixation, testes were washed 3 times in PBS and incubated in 30% sucrose, 0.02% sodium azide overnight at 4°C. Afterwards, testes were frozen in OCT (Sakura Finetek Europe) on dry ice and stored at -80°C until sectioned. 7 µM thick sections were cut with Leica CM1850, sections were dried on glass slides at room temperature. Slides were washed several times with distilled water to remove OCT, after which they were air dried. Dry slides were blocked with 2.5% w/v BSA in PBS for 1 hour, and then slides were stained as described for surface spreads.

To assess oocyte numbers in adult mice, DDX4 was detected in paraffin-embedded sections of ovaries in young adults (6 weeks old). Ovaries were dissected and fixed in 4% paraformaldehyde in 100 mM sodium phosphate buffer pH 7.4 overnight at 4C. Afterward, ovaries were washed 3 times in PBS pH 7.4, once with 70% ethanol and embedded in paraffin for sectioning at 5 mm thickness. Deparaffinization and rehydration of the sections was performed as follows: 2x 5 min in xylene, 2x 5 min in 100% ethanol, 5 min each in 95%, 85%, 70%, 50% ethanol, 2x 5 min in water. Sections were subjected to heat-mediated antigen retrieval in 10 mM Sodium citrate, 0.05% Tween 20, pH 6.0 for 20min in boiling water bath.

Sections were permeabilized in PBS with 0.2% Triton X-100 for 45 min at room temperature and processed for immunofluorescence staining immediately. DDX4-positive oocytes were counted on every seventh section of both ovaries in each female mouse.

### Staging of meiotic prophase in nuclear spreads

The first meiotic prophase substages were identified by a combination of three markers, SYCP3 (chromosome axis marker), SYCP1 (SC marker) and Histone H1t (post-mid pachytene marker in spermatocytes) (for details see also ^24^). We defined preleptotene as a stage where hazy/punctate staining pattern of SYCP3 is observed throughout the nucleus as described previously ^22^. The next stage, leptotene, is characterized by short stretches of axes and without any SC. This is a stage where recombination is initiated in wild type. The next stage is early zygotene, which is characterized by long but fragmented and incomplete axis stretches. In SC proficient genotypes, SC is also detected in this stage. Advancing to late zygotene stage, axes of all chromosomes are fully formed but SCs are incomplete. Cells enter pachytene when SC formation is completed in wild type. Whereas all chromosomes are fully synapse in pachytene oocytes, only autosomes synapse fully and heterologous sex chromosomes synapse only in their PARs in spermatocytes. SC formation was occasionally defective in *Iho1^C7Δ/C7Δ^*spermatocytes; hence we applied modified criteria for pachytene stage to include cells that have synaptic defects. Prophase stage was considered pachytene in spermatocytes if axes fully formed, axes had condensed appearance typical of pachytene, and the status of SC satisfied one of the following criteria: i) if PARs of sex chromosomes were synapsed ii) if all autosomes were fully synapsed but PARs of sex chromosomes were unsynapsed iii) if PAR asynapsis was accompanied with partial or full asynapsis on up to 3 autosome pairs. Histone H1t staining was used to sub-stage mid-or late-pachytene. Histone H1t staining is absent or weak in early pachytene but intermediate and high Histone H1t levels are seen in mid and late pachytene, respectively. In diplotene stage, axes desynapse and the SC becomes fragmented. Histone H1t levels remain high in this stage in spermatocytes. In oocytes, the same stages exist but due to lack of H1t, the developmental time of fetuses was used to aid staging as before ^64^. Most oocytes are in zygotene and mid-pachytene in fetuses 16 and 18 days postcoitum.

### Staging of mouse seminiferous tubule cross sections

Coordinated mitotic proliferation, meiosis entry and migration of spermatogenic cells to the lumen of seminiferous tubules result in 12 well-defined stages of the seminiferous epithelial cycle (I-XII) that are characterized by distinct associations of premeiotic, meiotic and postmeiotic spermatogenic cell layers across cross sections of seminiferous tubules ^94^.

We used DAPI in combination with histone-H1t (marker of spermatocytes after mid pachytene or stages V-XII) to stage the epithelial cycle as before ^24, 95^. Cleaved-PARP was used to label and quantify apoptotic spermatocytes.

### Testis organ culture

Organ cultures of testes were carried out as described earlier ^34, 96^ with minor modifications. Testes of 8 dpp mice were dissected, tunica albuginea was removed and each testis of juvenile mouse was split to 2 or 3 pieces. Freshly isolated tubules were cultured at gas/liquid interphase on agarose gel blocks (1.5%; W/V; thickness about 7mm). Gel pieces were preincubated in culture medium for 24 hours to saturate agarose with medium before samples were placed on them. After the biological samples were placed on the agarose slices, the medium level was adjusted not to cover the seminiferous tubules in the culture wells. We used α-MEM (Life Technologies), 10% (v/v) KSR (Life Technologies) and gentamycin (Sigma-Aldrich) at a final concentration of 5 μg/ml as culture medium. The culture medium was supplemented with CDC7 inhibitors at concentrations indicated in the figure legends. We used various combinations of three inhibitors: TAK-931, XL413 hydrochloride and LY3143921 hydrate. Stock solutions of inhibitors were prepared per manufacturer’s instructions. Tubules were incubated at 34°C in humidified atmosphere containing 5% CO2. After 2 days of incubation, samples were either processed for nuclear surface spreads or frozen for WB analysis.

### Multiple sequence alignments

Multiple sequence alignments was done using Clustal Omega from the webserver of EMBL-EBI^97^. Jalview was used to edit alignments ^98^.

### Image quantification

Fiji variant of ImageJ was used to manually quantify Western blot band intensities ^99–101^. Background of each lane was subtracted and signal was normalized to loading control of the same lane.

To reliably detect functional DSB-machinery clusters, we identified MEI4 foci (guinea pig-MEI4 antibodies) that colocalized with REC11 foci (rabbit-REC114 antibodies) ^22^. Focus numbers of DSB factors (MEI4 and REC114) were quantified by Cell Profiler 3.0 software ^102^. Macros of the Cell Profiler anylasis are accessible at https://github.com/IhsanDereli/Iho1_C7del. A difference of gaussian blurs was used to define the borders of each MEI4 and REC114 focus similar to a previously used approach for the identification of DSB-factor foci ^23^ except that instead of automatic thresholding we empirically selected thresholds that allowed correct identification of DSB-clusters according to manual inspection in all of the examined meiotic stages for each experimental set. We considered a MEI4 focus as a functional DSB-machinery cluster only if at least 50% of the area of the MEI4 focus overlapped with a REC114 focus. The original grayscale images were masked based on identified foci and integrated signal intensity of MEI4 was calculated on original unprocessed images, after corrected for varying background signal by subtracting median of all pixel intensity values in the image.

ssDNA associated protein foci were counted manually.

ImageJ was used to process images for DMC1-RPA2 density measurements. First, we identified nuclear spreads where it was possible to delineate individual axis and SC stretches. Synapsis was identified by the presence of SYCP1 or depletion of HORMAD1. ‘Segmented line’ tool was used to mark and measure the length of synapsed and unsynapsed regions. DMC1 and RPA2 focus numbers were manually counted on synapsed or unsynapsed regions and DMC1 and RPA2 focus numbers were added up to estimate recombination focus numbers. If a DMC1 and an RPA2 focus overlapped or touched each other, we considered them as a single recombination focus if the center of one of the foci overlapped with the signal of the other focus. Recombination focus densities were calculated by dividing the sum of DMC1 and RPA2 foci in synapsed or unsynapsed regions with the total length of synapsed or unsynapsed chromosome axis, respectively, in each cell.

### Quantification and statistical analysis

Graphs were plotted by Graphpad Prism 9. All statistical tests were done using R version 4.1.3. The statistical analyses were implemented in the R packages lmerTest and lme4 ^103, 104^. The R code was published previously ^95^.

### Data and software availability

The mass spectrometry proteomics data have been deposited to the ProteomeXchange Consortium via the PRIDE ^105^ partner repository with the dataset identifiers PXD042179 and PXD042221 and 10.6019/PXD042221.

---

**Figure S1.**
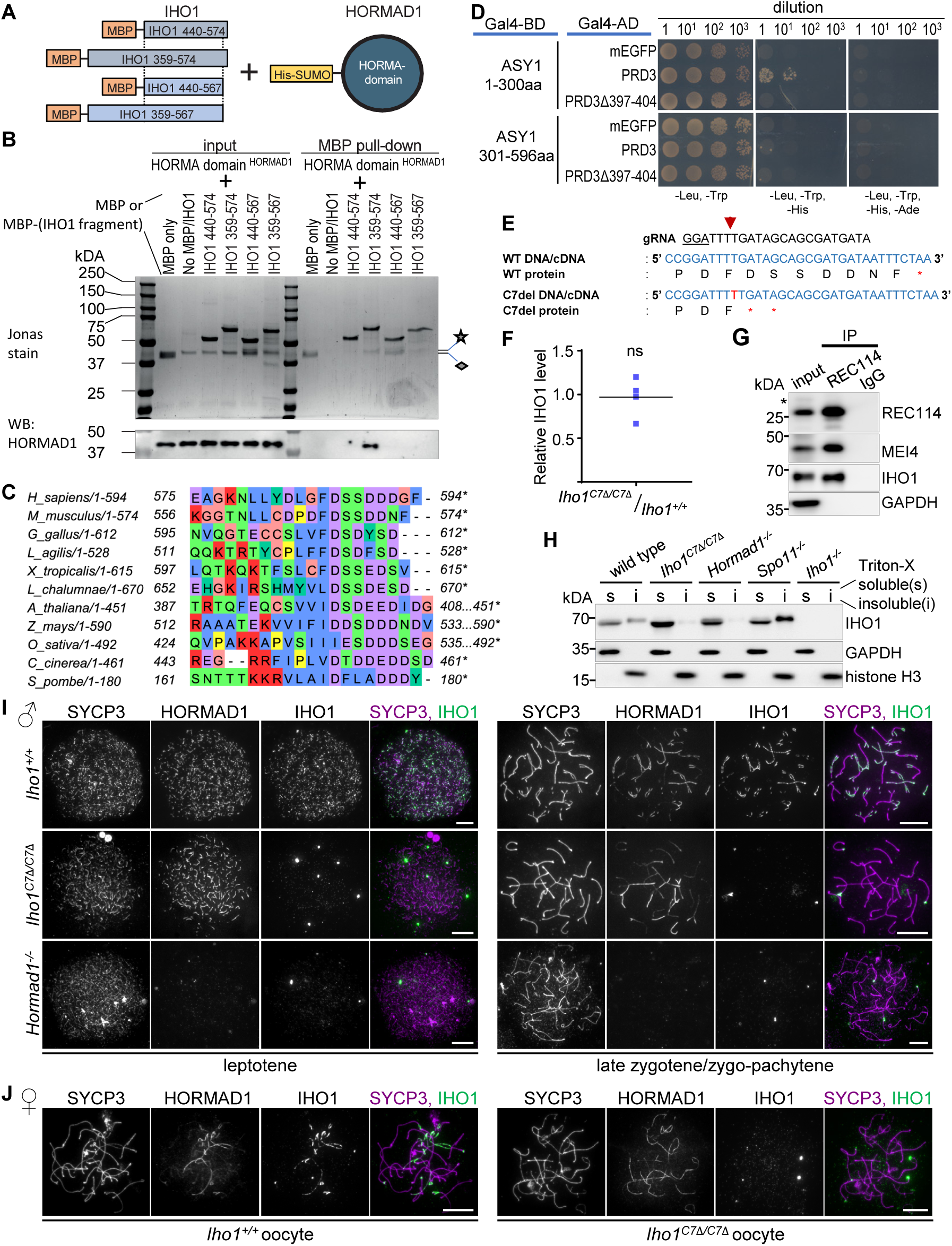
A conserved motif in IHO1 C-terminus is required for axis localization of IHO1 in both sexes (relevant to Fig. 1) **A** Schematics of *in vitro* interaction assays between HORMA domain of HORMAD1 (1-240 amino acids) and fragments of IHO1 (the amino acid position of fragment boundaries are indicated). **B** Jonas stained gel picture (top) and immunoblot analysis (WB, bottom image) of *in vitro* MBP pull down assays testing interaction between purified His6-SUMO-tagged version of the HORMA domain of HORMAD1 (1-240 amino acids) and MBP-tagged versions of IHO1 fragments as depicted in **A**. Star and diamond mark the nearly indistinguishable positions of the protein bands of His6-SUMO-HORMAD1 HORMA domain (40.3kDa) and MBP, respectively. Note that the ∼40kDa protein band that is detected in the MBP-IHO1 440-567 pull down by Jonas staining is probably a degradation product of MBP-IHO1 440-567 as it is not detected by the HORMAD1 immunoblot. **C** A window of a multiple sequence alignment of the last 100 amino acids of Mer2/IHO1-family proteins in diverse taxa, showing the previously described short similarity motif/SSM2 ^14^ and an overlapping/adjacent conserved acidic patch at its C-terminus. Amino acid positions of the beginning and the end of the window and the positions of the last amino acids (*) are shown in full length proteins. **D** Yeast two-hybrid interaction assays between *Arabidopsis thaliana* proteins. Interactions were tested between the N-terminal 300 amino acids of ASY1 (HORMAD1 orthologue) including the HORMA domain or a C-terminal ASY1 fragment (amino acid positions 301-596) containing a SWIRM domain and either wild-type PRD3 (IHO1 orthologue) or a PRD3 version that lacks the conserved IDSDEED motif (PRD3Δ397-404). Y2H assays between ASY1 fragments and mGFP served as negative controls. Cell suspensions of indicated optical densities (OD) were plated, and yeast cultures are shown after 6 days of growth on dropout plates. **E** Genomic DNA and corresponding protein sequence of IHO1-Cterminus in wild-type (WT) and *Iho1^C7Δ/C7Δ^* mutant line (C7del). Guide RNA sequence (gRNA) used for CRISPR/Cas9 editing is shown. PAM (underlined) and DNA cut site (red arrow) are marked. Red text color identifies mutated DNA and protein sequence, * marks STOP codons in the edited locus. **F** Quantification of total IHO1 protein levels in *Iho1^C7Δ/C7Δ^* mice. Immunoblot signals of total IHO1 protein from testis extracts of 13 dpp *Iho1^C7Δ/C7Δ^*mice were normalized to corresponding immunoblot signals of 13 dpp wild-type controls. Bar marks mean=0.97 of four experiments; two-tailed one-sample *t* test, ns=0.8055. **G** Immunoblots of immunoprecipitations (IP) from testis extracts of wild type mice at 13 dpp age. Immunoprecipitations with non-specific rabbit immunoglobulins (IgG) served as a negative control for IPs with rabbit anti-REC114 antibodies. Asterisk mark aspecific band that appears with variable intensity (compare to Fig. 1E) in REC114 immunoblots of IP-input testis extracts. **H** Immunoblot analysis of Triton X (0.3%) soluble (s) and insoluble (i) fractions of extracts of testes from 13 dpp mice of indicated genotypes. Panel shows IHO1, GAPDH (a marker of soluble fraction) and histone H3 (a marker of insoluble chromatin fraction). **I-J** Immunofluorescence of IHO1, and markers of the chromosome axis (SYCP3) and unsynapsed axis (HORMAD1) in nuclear surface spread spermatocytes (**I**) of adult mice in indicated stages or late zygotene oocytes (**J**) of 16.5 days postcoitum (dpc) fetuses. Bars, 10 µm.

**Figure S2.**
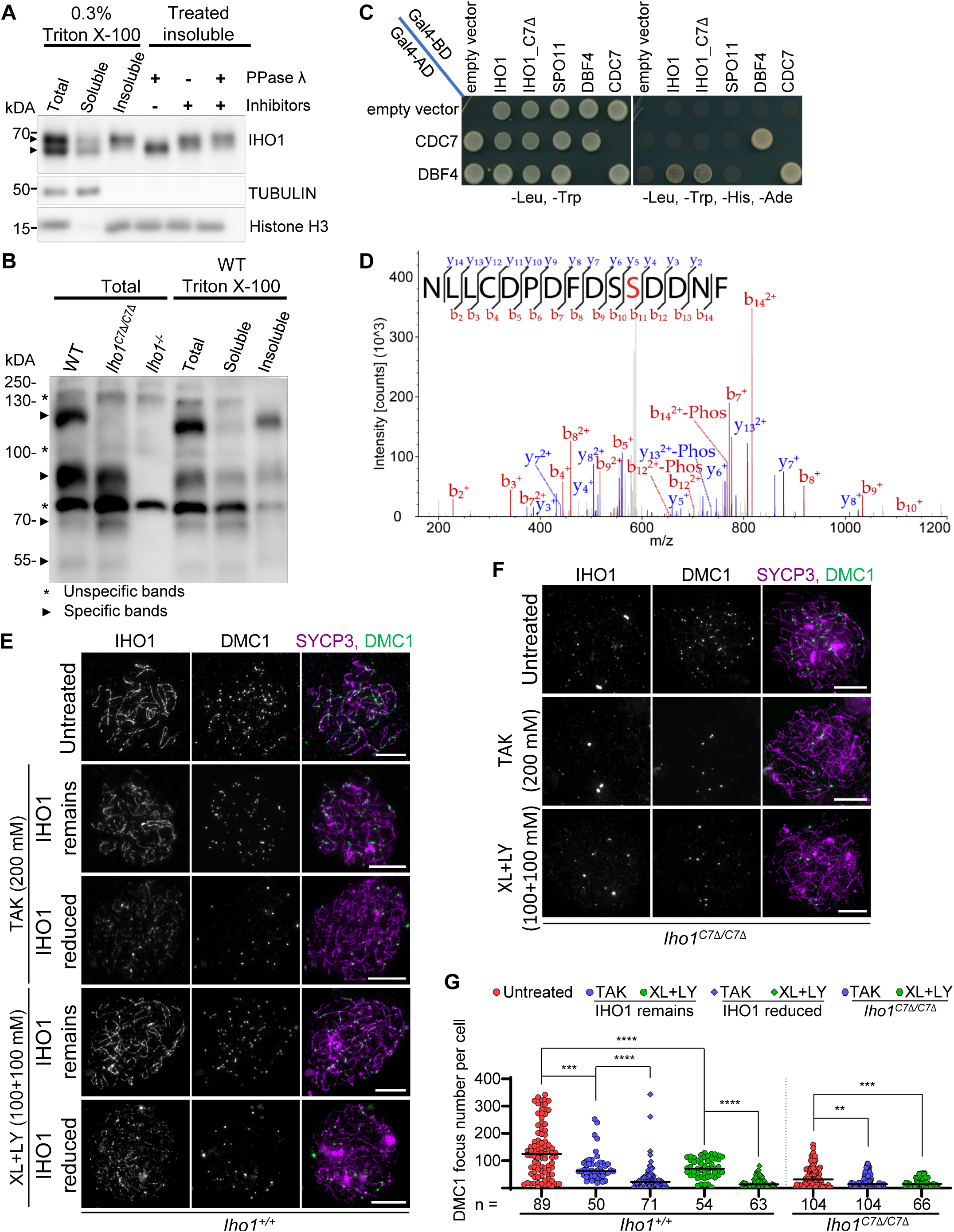
Analyses of IHO1 phosphorylation, interactions between IHO1 and CDC7/DBF4, and phenotypes of CDC7 inhibition in meiosis (relevant to Fig. 2) **A, B** Immunoblots of standard (**A**) or phos-tag (**B**) SDS-PAGE of protein extracts from testes of 13 dpp mice. **A** Fractionation of testis extracts based on Triton X-100 solubility (left 3 lanes) and λ phosphatase treatment of Triton X 100-insoluble fraction (right 3 lanes) of wild-type testes. **B** Total testis extracts of mice with indicated genotypes (left 3 lanes) and fractionation of testis extracts of wild-type (WT, right 3 lanes) mice. Unspecific bands were marked by asterisk. Black triangles mark four distinct IHO1 specific bands that indicate an unphosphorylated form plus at least three phosphorylated forms for wild-type IHO1 and an unphosphorylated form plus at least two residual phosphorylated forms for IHO1_C7Δ. **C** Yeast two-hybrid interaction assays between indicated proteins. Yeast cultures are shown after 3 days of growth on dropout plates. For negative control, proteins of interest were tested in transformations where either the Gal4-binding domain (Gal4-BD) or the Gal4-activation domain (Gal4-AD) vectors were empty. **D** MS/MS spectrum of NLLCDPDFDS(pS)DNF-NH_2_ peptide. Identified b and y ions are annotated in red and blue, respectively. Fragments with neutral loss of H_3_PO_4_ are indicated as “-Phos”. 9+/-1.4% of the peptide was phosphorylated in n=3 measurements. **E-G** Analysis of spermatocytes from testes of 8 dpp mice after 48 hours *in vitro* culture with or without CDC7 inhibitors 200mM TAK-931 (TAK) or a mix of 100mM XL413 and 100mM LY3143921 (XL+LY). **E-F** Immunofluorescence of IHO1, markers of the chromosome axis (SYCP3) and unrepaired DSBs (DMC1) in nuclear surface spread spermatocytes of *Iho1^+/+^* (**E**) or *Iho1^C7Δ/C7Δ^* (**F**) mice. **E** Spermatocytes are shown with normal or reduced levels of axial IHO1 in CDC7 inhibitor treated *Iho1^+/+^* samples. **G** Quantification of DMC1 foci, datapoints represent cells from 2 pooled experiments. DMC1 quantifications are shown separately for *Iho1^+/+^* cell populations where IHO1 remained on axes and cell populations where IHO1 was depleted from axes following treatment with CDC7 inhibitors. Bars, medians. Mann-Whitney U test, s**=P<0.01, s***=P<0.001, s****=P<0.0001.

**Figure S3.**
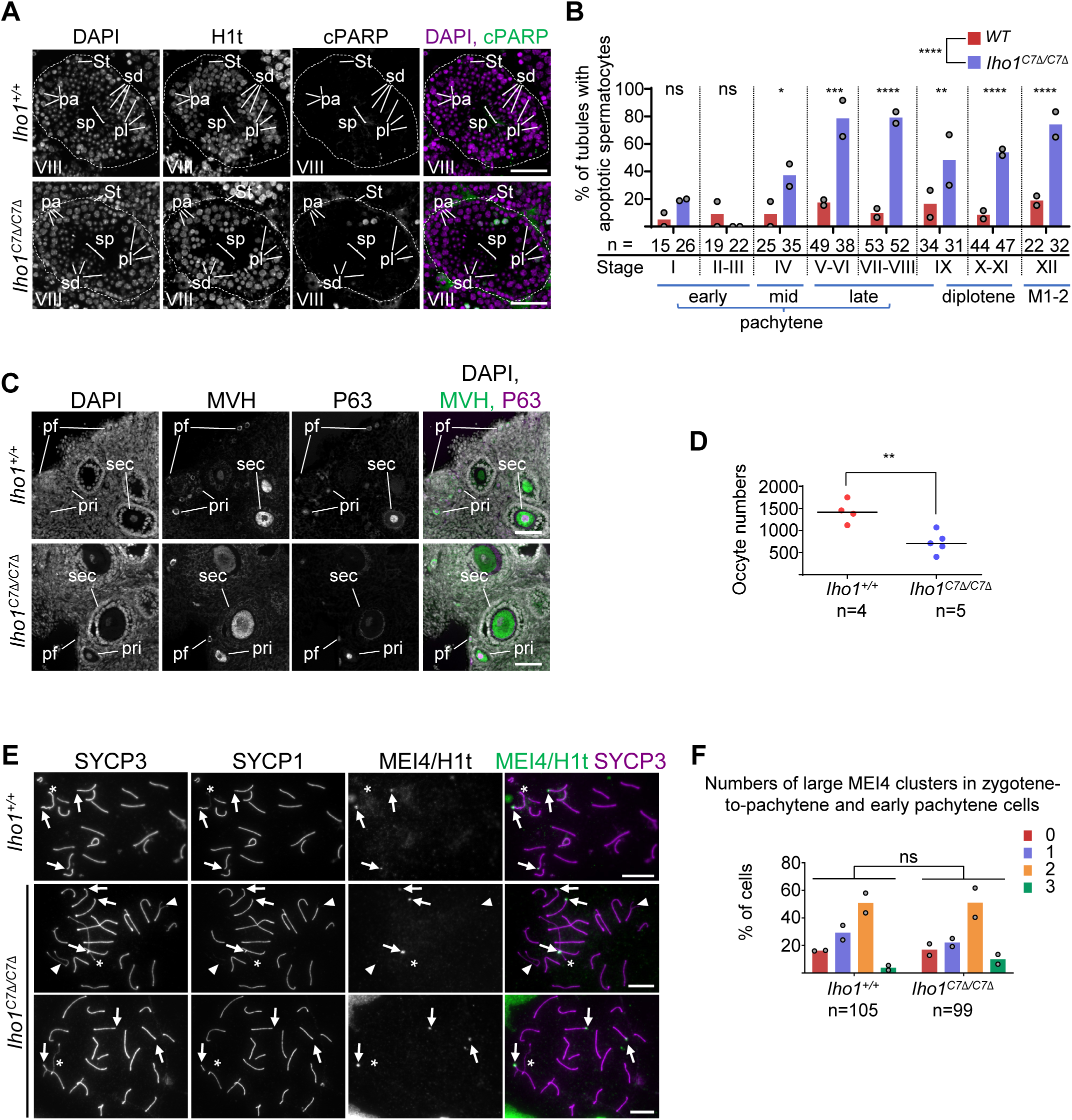
*Iho1^C7Δ/C7Δ^* mice show increased loss of meiocytes but no effect on laDSB clusters (relevant to Fig. 3) **A** Sections of testes from adult *Iho1^+/+^* and *Iho1^C7Δ/C7Δ^* mice. DNA was detected by DAPI. Histone H1t (staging marker of seminiferous tubules) and cleaved PARP (cPARP, marker of apoptosis) were detected by immunostaining. Epithelial cycle stages of the seminiferous tubules are indicated by roman numbers. Preleptotene (pl), pachytene (pa), round spermatid (sd), sperm (sp) and Sertoli cells (St) and outlines of tubules (white dashed line) are marked. **B** Quantification of seminiferous tubules that contain cPARP-positive apoptotic spermatocytes in the pachytene/diplotene/mitotically dividing cell layers. Block bars show the averages from two experiments, gray circles represent single experiments. n = number of tubules counted in two independent experiments. Analysis of deviance using the likelihood-ratio test based on the chi-squared distribution, ns=P>0.05, s*=P<0.05, s**=P<0.01, s***=P<0.001, s****=P<0.0001. Most advanced meiotic cell-division cycle stage is indicated in the epithelial cycle stages. **C** Cryo sections of ovaries from young adult mice (6 weeks). Two oocyte markers, MVH (cytoplasmic) and p63 (nuclear) were immunostained, DNA was labeled by DAPI. Primordial (pf), primary (pri) and secondary (sec) follicles are indicated. **D** Quantification of oocyte numbers in ovary sections from 6-weeks-old females of indicated genotypes. Sums of oocyte numbers from every 6th sections of both ovaries of each mouse are shown. *n* = numbers of analyzed animals; two tailed Welch t-test, s**=P<0.01. Bars mark mean numbers of oocytes (wild-type, 1424.25, *Iho1^C7Δ/C7Δ^*, 728). **E** Immunostaining of surface spread spermatocytes of adult mice. MEI4 was detected in the same channel as histone H1t, as these two proteins mark distinct populations of spermatocytes, up till early pachytene and beyond mid pachytene, respectively. Panels show correctly synapsed early pachytene cells (*Iho1^+/+^* and *Iho1^C7Δ/C7Δ^*bottom panel) and an *Iho1^C7Δ/C7Δ^* cell with incomplete synapsis (middle panel) in zygotene-to-pachytene transition or early pachytene-equivalent stage. Arrows, asterisk and arrowheads mark large MEI4 clusters, sex chromosomes, asynaptic autosomes, respectively. **F** Quantification of MEI4 blob numbers in zygotene-to-pachytene transition and early pachytene spermatocytes of adult mice. Block bars show averages of two experiments, gray circles represent single experiments. Total numbers of counted cells are indicated. Analysis of deviance using the likelihood-ratio test based on the chi-squared distribution, ns=P>0.05. **A, C, E** Bars, 100µm (**A**), 20 µm (**C**), 10 µm (**E**).

**Figure S4.**
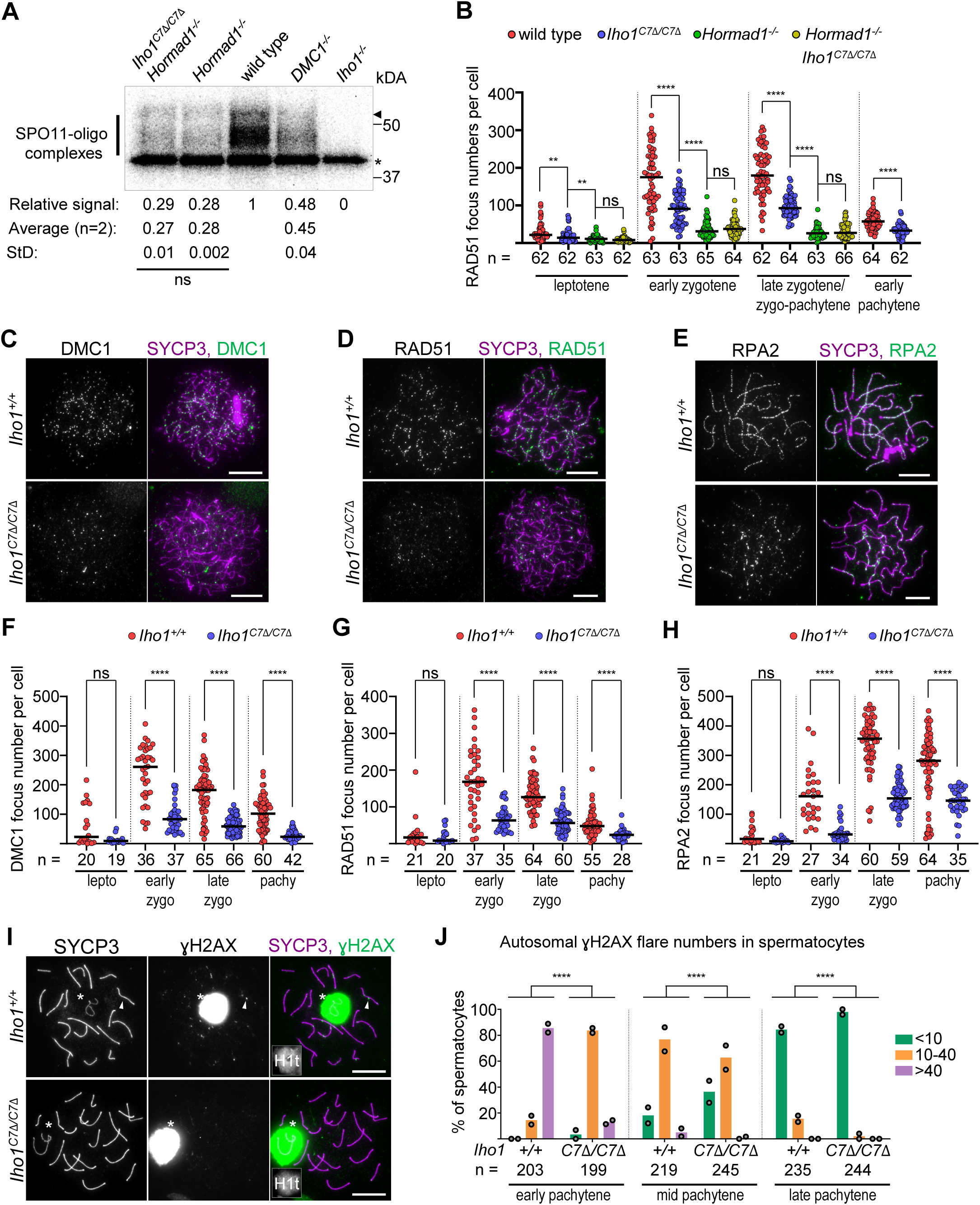
Disruption of IHO1-HORMAD1 interaction affects early recombination steps (relevant to Fig. 4) **A** Radiograph of immunoprecipitated and radioactively labeled SPO11-oligo complexes from testes of adult mice. Bar, SPO11-specific signals, asterisk, nonspecific labelling, and arrowhead, immunoglobulin heavy-chain. Quantification shows radioactive signals that were background-corrected (*Iho1^-/-^*, signal=0) and normalized to wild type control (1). Means and standard deviations of SPO11-oligo signals are shown from n=2 independent experiments (radiograph shows one of the experiments). Paired t-test, ns=P>0.05. **B** Quantification of axis associated RAD51 focus numbers in spermatocytes. Bars are means, n=cell numbers. Mann Whitney U-test, ns=P>0.05, **=P<0.01, ****=P<0.0001. **C-E, I** Immunostaining of nuclear spread oocytes of 16.5 days-post-coitum (dpc) fetuses (**C-E**) or nuclear spread spermatocytes of adult mice (**I**). Bars, 10 µm. **C-E** Oocytes are shown in early zygotene (**C-D**) or late zygotene (**E**) stages. **I** Spermatocytes are shown in late pachytene stage. Miniaturized images show histone H1t. Sex chromosomes are marked by asterisk. Arrowhead in top panel points at a ɣH2AX flare/foci on a synapsed autosome. **F-H** Quantification of axis associated DMC1 (**F**), RAD51 (**G**) RPA2 (**H**) focus numbers in leptotene (lepto), early zygotene (early zygo), late zygotene (late zygo) and early pachytene (pachy) oocytes of 16.5 dpc fetuses. Bars are medians, n=cell numbers. Mann Whitney U-test, ns=P>0.05, ****=P<0.0001. **J** Quantification of autosomal ɣH2AX flare numbers in spermatocytes of adult mice. Three categories were distinguished, spermatocytes with <10 flares (green), between 10 and 40 flares (orange) and more than 40 flares (lilac) of ɣH2AX on autosomes. Whereas cells with asynaptic chromosomes were also included the quantifications, asynaptic chromosomes exhibit large ɣH2AX domains that are associated with the sex bodies, hence asynaptic chromosomes are not reflected in flare counts. Block bars show averages of two experiments, gray circles represent single experiments, n=number of counted cells. Analysis of deviance using the likelihood-ratio test based on the chi-squared distribution, ****=P<0.0001.

**Figure S5.**
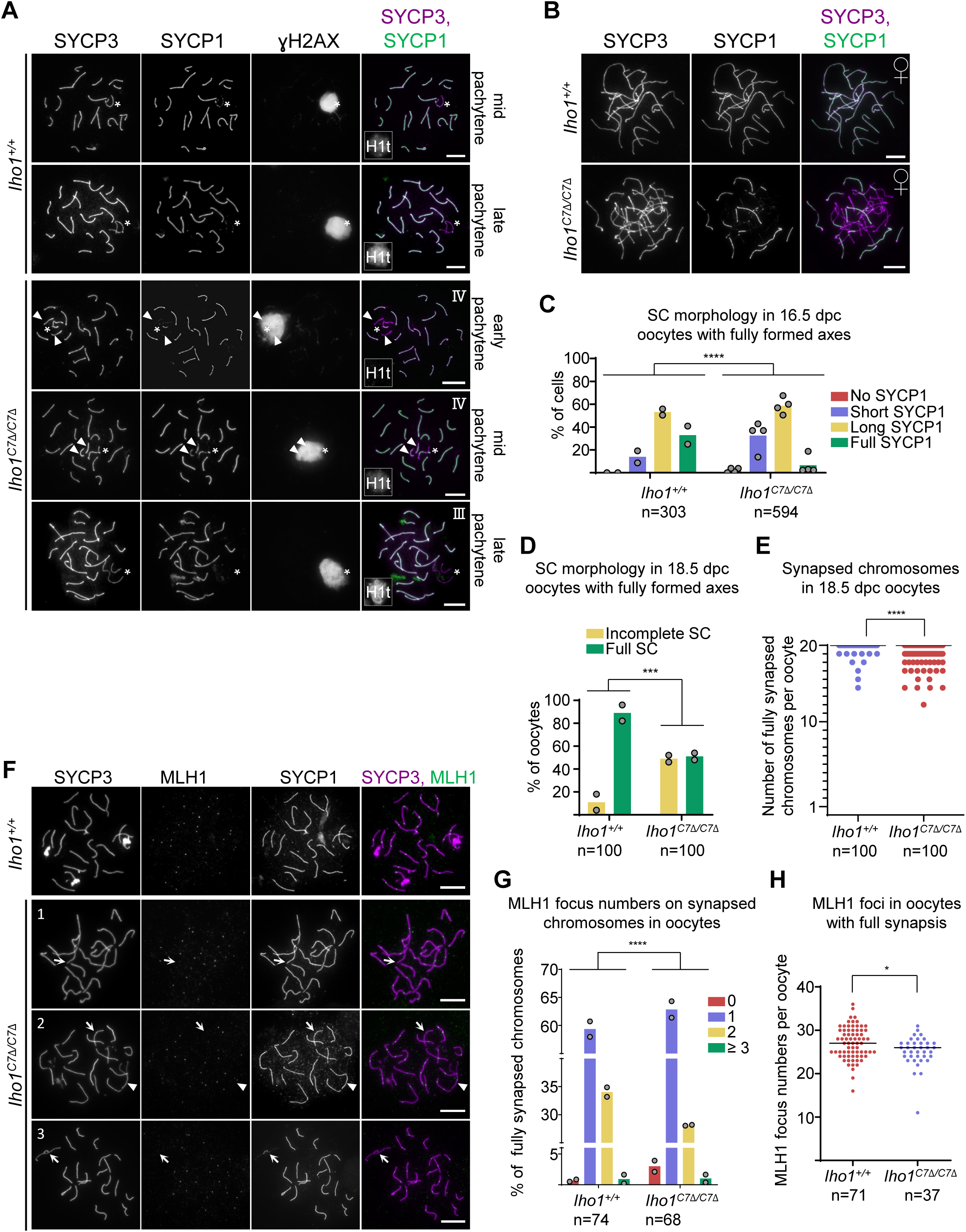
SC defects and imperfect CO formation affect both sexes in *Iho1^C7Δ/C7Δ^* mice (relevant to Fig. 5) **A-B , F** Immunostaining in nuclear spread spermatocytes of adult mice (**A**) or oocytes of 16.5 (**B**) or 18.5 (**C**) dpc fetuses. Bars, 10 µm. **A** Asterisks mark sex chromosomes. Arrowheads mark partially or fully unsynapsed autosomes. Miniaturized images show histone H1t. Roman numbers refer to SC defect types as described in Fig. 5B. **F** Three types of *Iho1^C7Δ/C7Δ^* oocytes are shown: (cell 1) all chromosomes are fully synapsed but one chromosome lacks MLH1 (arrow), (cell 2) one chromosome is partially synapsed and have MLH1 focus (arrowhead) and another fully synapsed chromosome lacks MLH1 (arrow), and (cell 3) all synapsed chromosomes have MLH1 foci but one chromosome that is partially synapsed lacks MLH1 focus (arrow). **C-E, G-H** Quantifications of SC morphology in oocytes with fully formed axis at 16.5 dpc (**C**) or 18.5 dpc (**D**), quantification of fully synapsed chromosomes in oocytes from 18.5 dpc fetuses (**E**), and quantifications of MLH1 focus numbers on fully synapsed chromosomes (**G**) or MLH1 focus numbers per cell (**H**) in oocytes of 18.5dpc fetuses. Both fully synapsed and asynaptic (**G**) or only fully synapsed (**H**) pachytene oocytes were examined. Data are pooled from two (**C,** *Iho1^+/+^*, **D-E**, **G-H**) or four (**C,** *Iho1^C7Δ/C7Δ^*) mice. Block bars are means (**C-D, G**), and bars are medians (**E, H**), n=numbers of oocytes. Likelihood ratio test (**C-D, G**) or Mann Whitney U-Test (**E, H**), *=P<0.05, ***=P<0.001, ****=P<0.0001.

**Figure S6.**
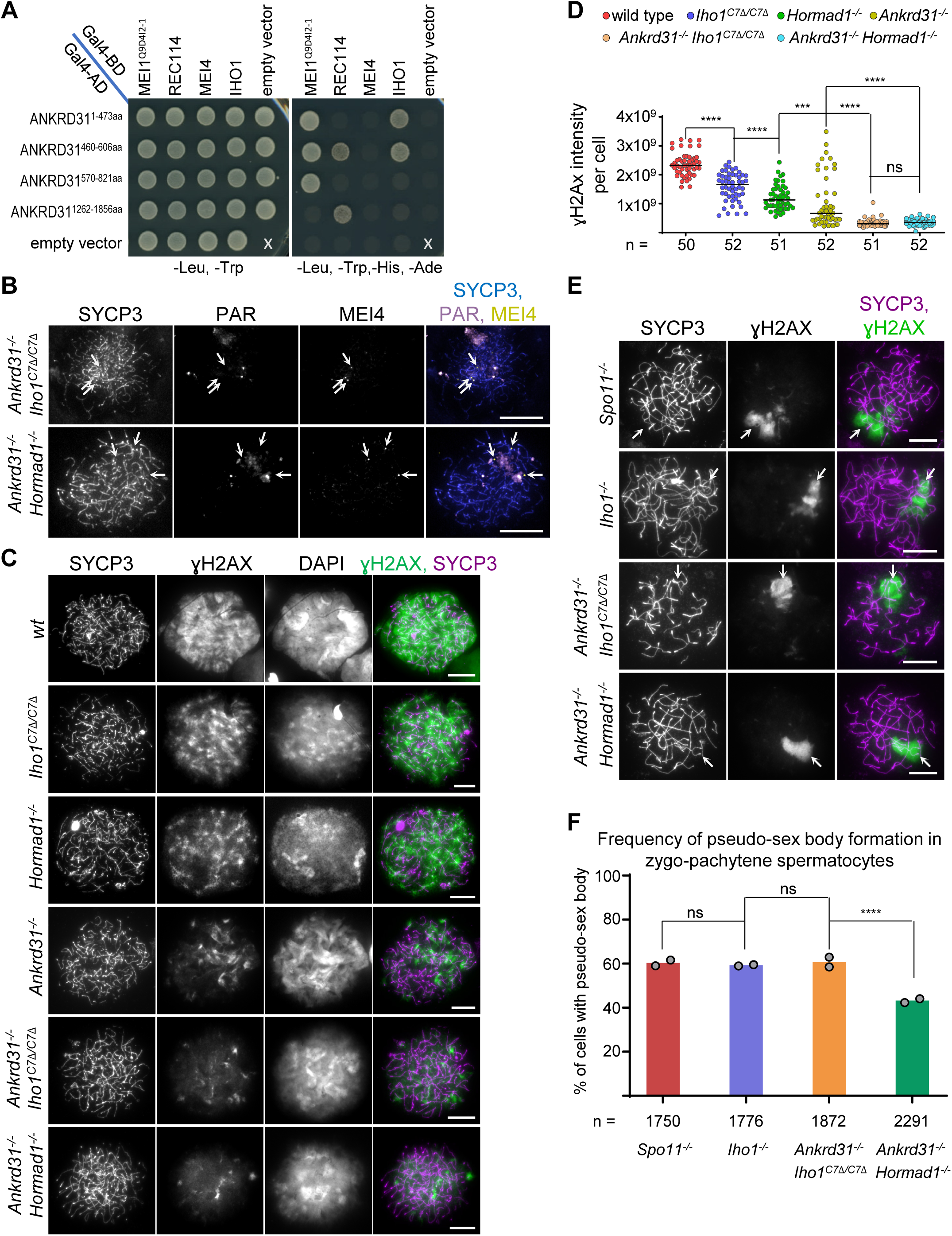
Physical and functional interactions of IHO1-HORMAD1 complex and ANKRD31 (relevant to Fig. 7) **A** Y2H assays between ANKRD31 fragments and key components of the DSB machinery. Yeast cultures are shown after 2 days of growth on the indicated drop out plates. **B-C, E** Immunostaining (**B-C, E**) and PAR FISH (**B**) in nuclear spread early zygotene (**B-C**, long but incomplete axes) or zygo-pachytene (**E**, fully formed axes) spermatocytes of adult mice. Arrows mark sites of MEI4 clusters (**B**) or ɣH2AX-rich chromatin domains, called pseudo-sex bodies (**E**). Bars, 10 µm. **D, F** Quantification of total nuclear ɣH2AX signal (in arbitrary units) in early zygotene spermatocytes (**D**) and quantification of the fraction of zygo-pachytene cells with pseudo-sex body formation (**F**) in adult mice. Bars are medians (**D**), block bars are means (**F**). Datapoints represent cells (**D**) or experiments (**F**). n=number of spermatocytes. Mann Whitney U-Test (**D**) or likelihood ratio test (**F**), ns=P>0.05, ***=P<0.001, ****=P<0.0001.

**Figure S7.**
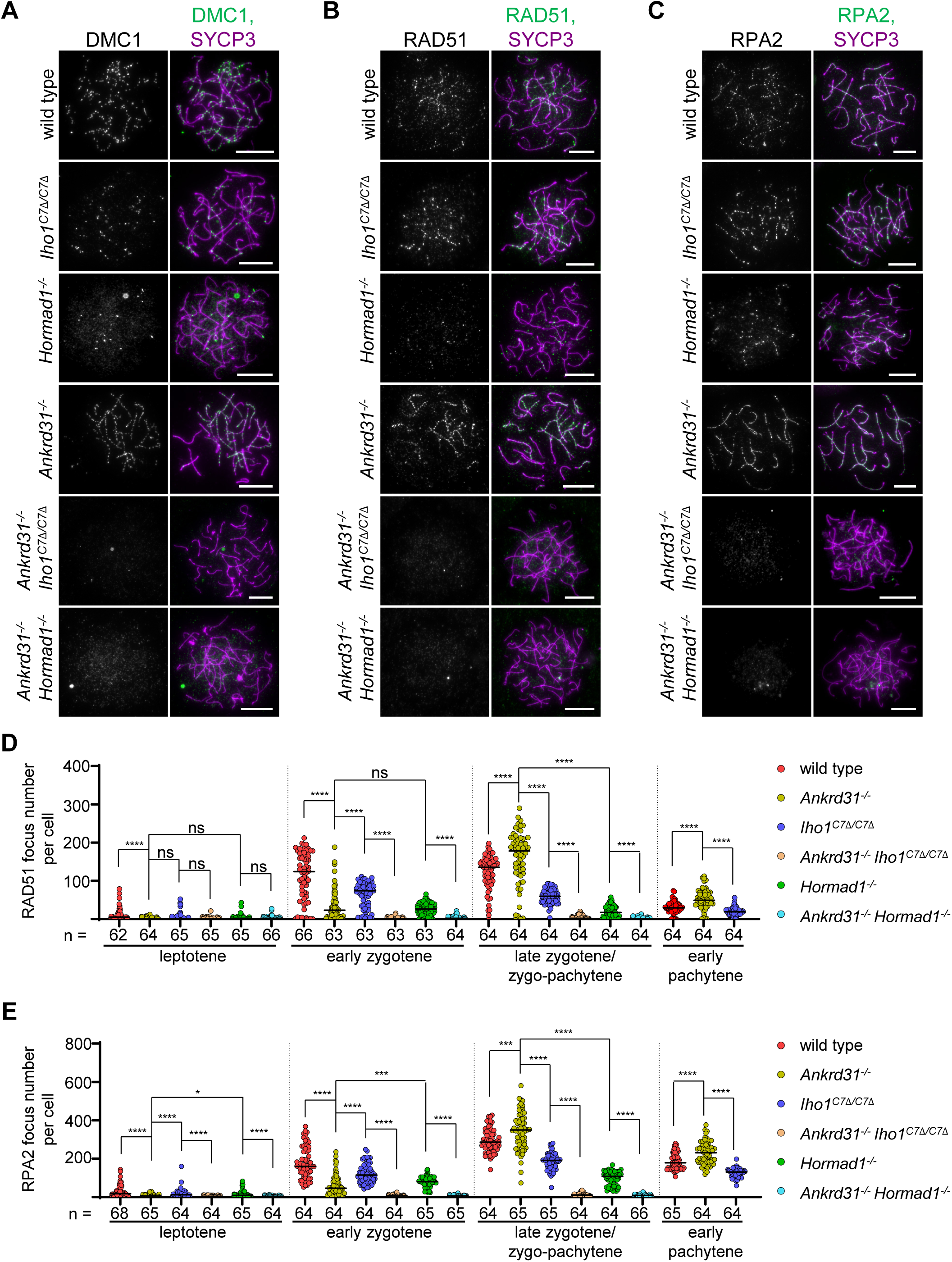
Recombination foci are abolished when both IHO1-HORMAD1 complex and ANKRD31 are lost (relevant to Fig. 7) **A-C** Immunostaining in nuclear spread late zygotene spermatocytes of adult mice. Bars, 10 µm. **D-E** Quantifications of RAD51 (**D**) and RPA2 (**E**) focus numbers in spermatocytes of adult mice. Zygo-pachytene (relevant only to severely SC-defective backgrounds, *Ankrd31^-/-^ Iho1^C7Δ/C7Δ^*, *Hormad1^-/-^*, *Ankrd31^-/-^ Hormad1^-/-^*) is equivalent to a mix of late-zygotene and early pachytene stages which are indistinguishable if SC is defective. Bars are medians, n=cell numbers. Mann Whitney U-Test, ns=P>0.05, *=P< 0.05, ***=P<0.001, ****=P<0.0001.

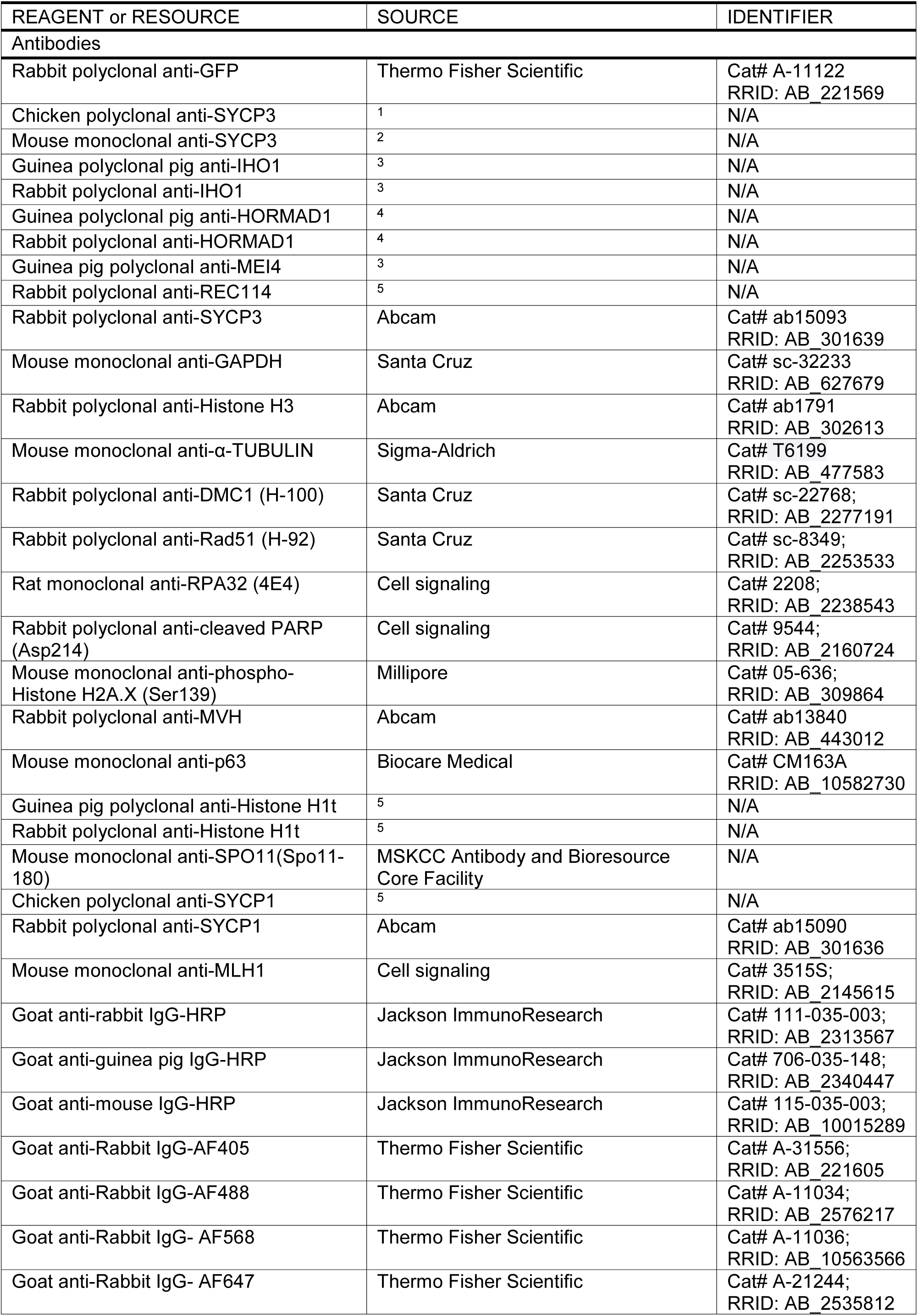
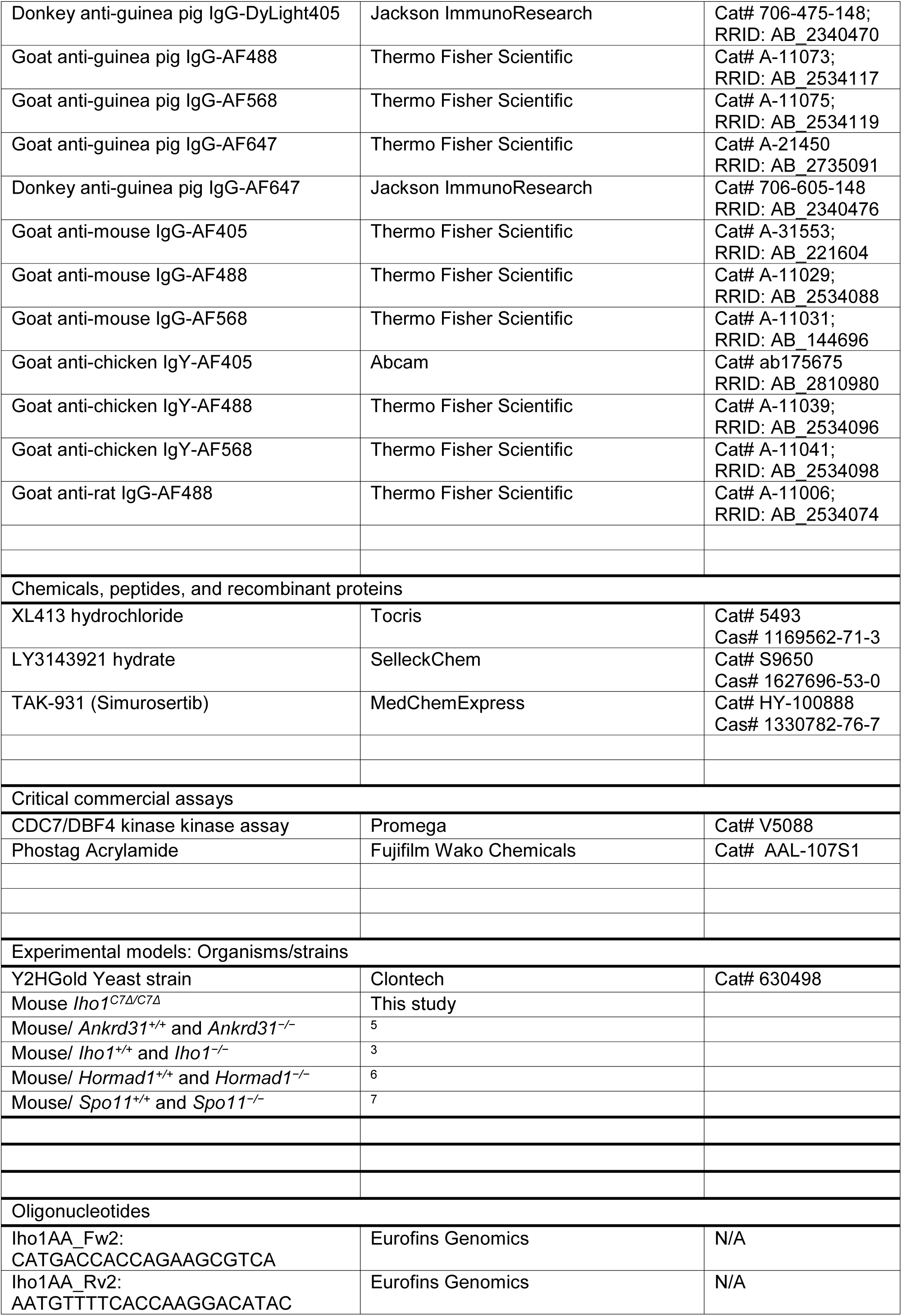
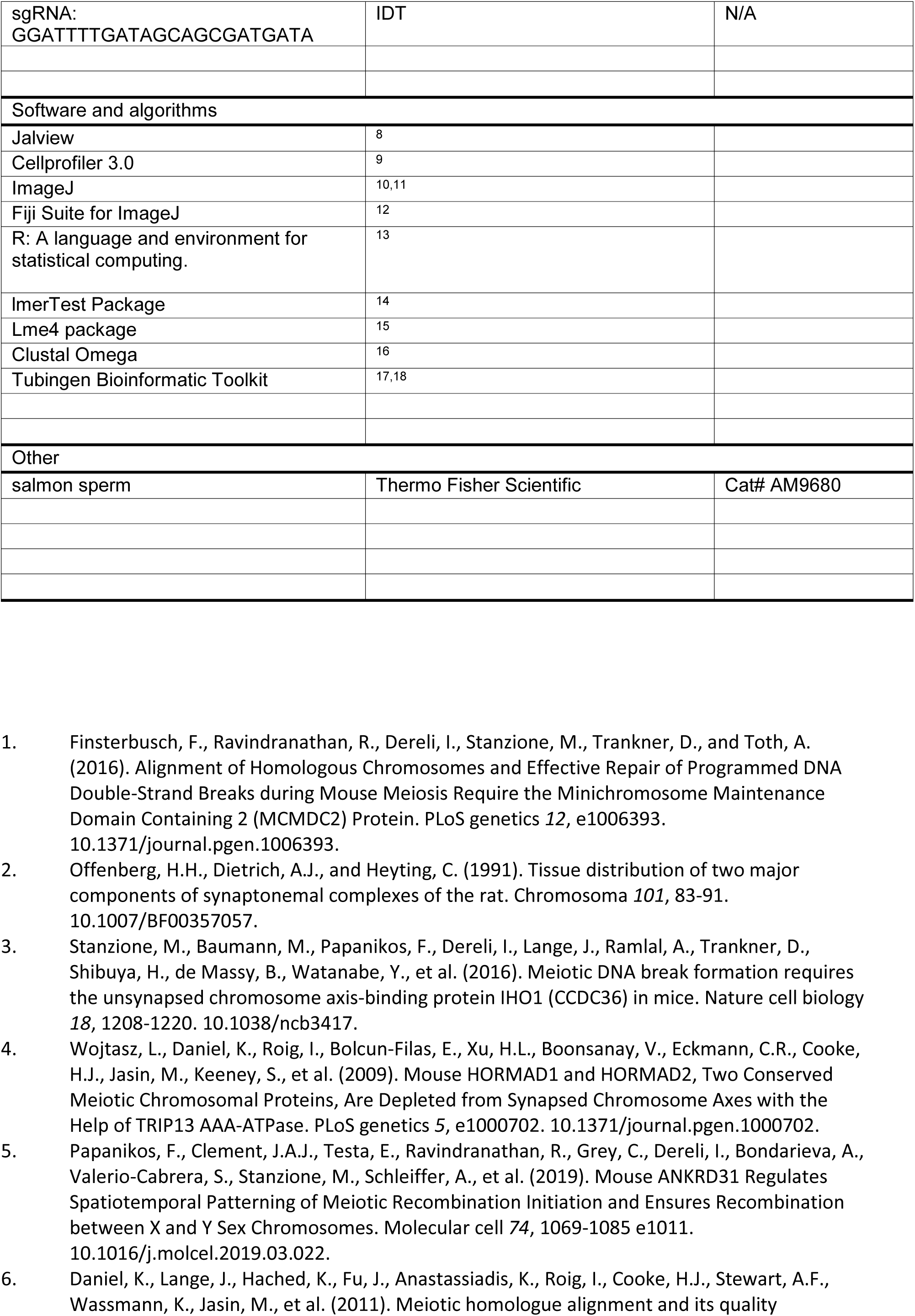
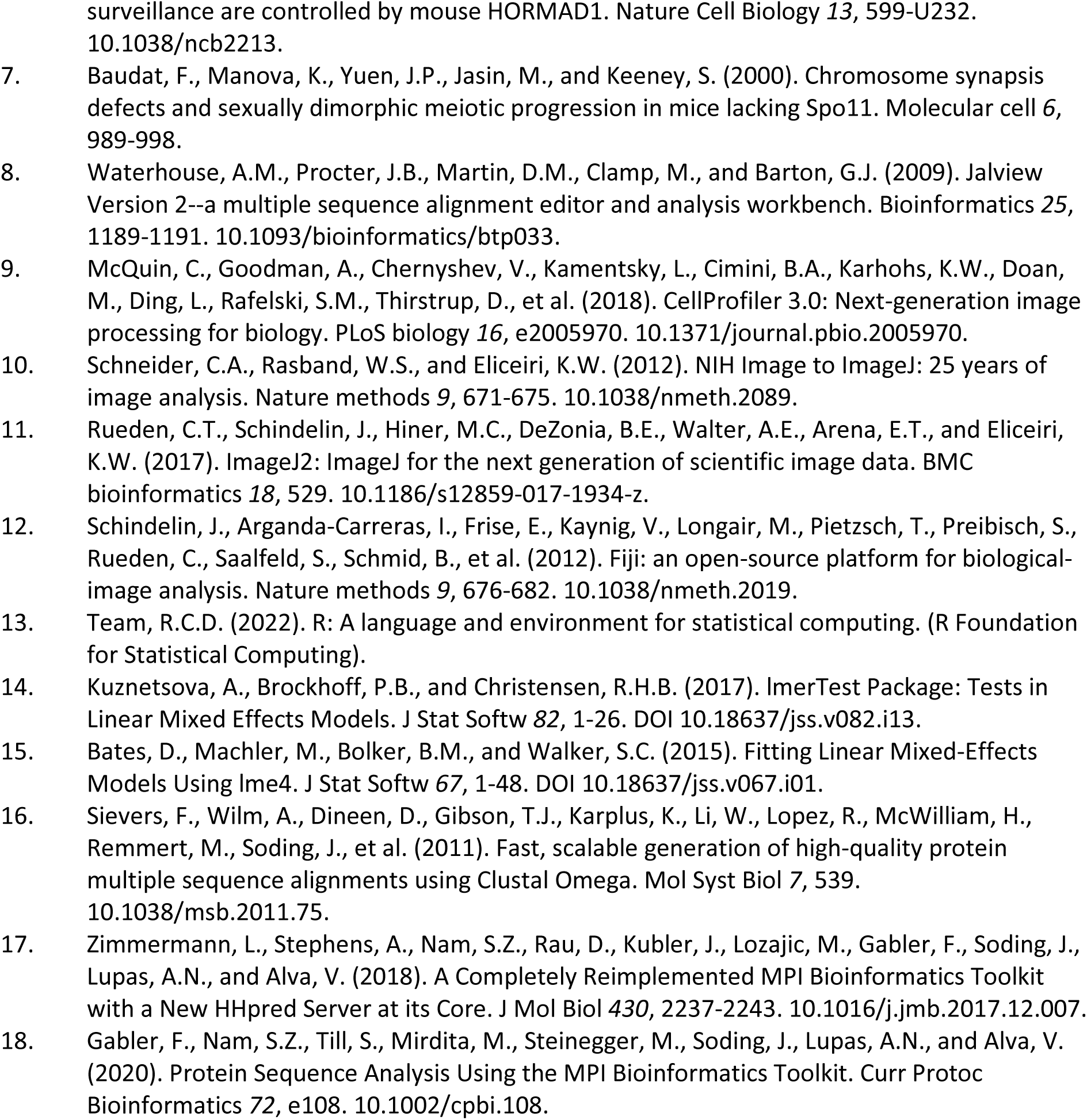
Table of reagents and resources.

**Table S1.**
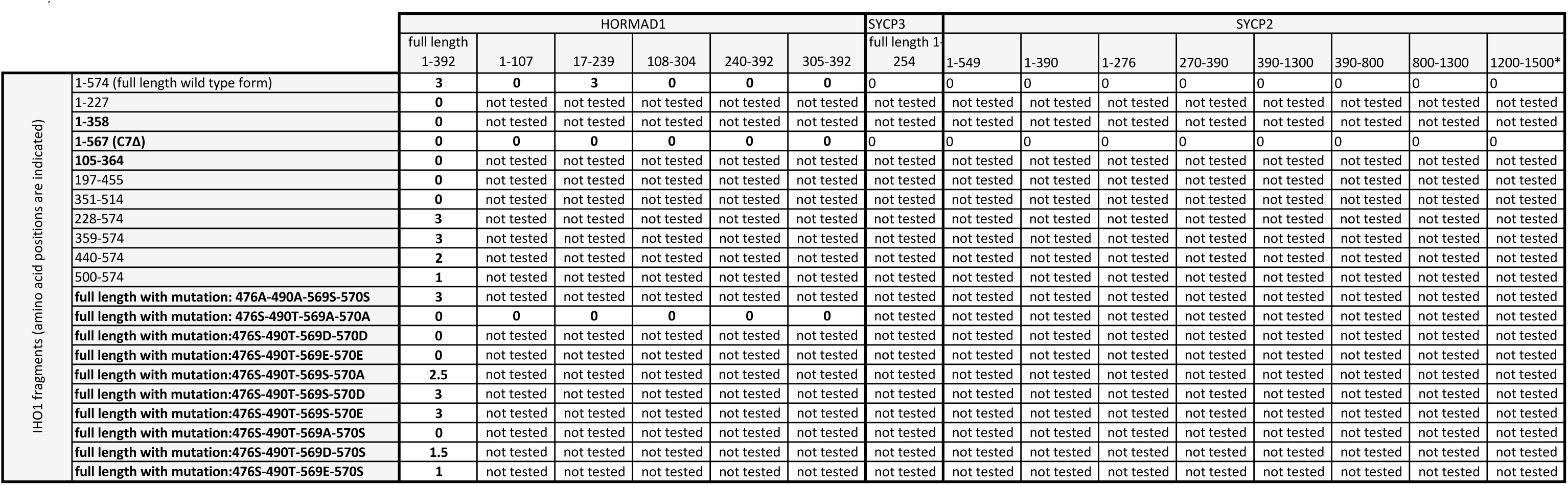
Y2H assays between full length and diverse versions of IHO1 (Gal4AD) and axis components HORMAD1, SYCP2 or SYCP3 (Gal4BD) (relevant to Fig. 1) Growth of budding yeast in Y2H assay (-Leu, -Trp, -His, -Ade plates) was graded between 0 (no growth) and 3 (strong growth). Grading is not directly comparable in tables S1 and S6. * SYCP2 fragment (1200-1500)-GAL4BD autoactivates leading to growth after three days; 0 grading refers to no additional growth relative to autoactivation in this case. Amino acid positions of the beginning and the end of protein fragments are indicated. Bold fragments of IHO1 contained a coiled coil domain, as predicted by alphafold (amino acid positions 114-246). In mutated forms of IHO1, we indicate the positions and mutations of the serines and the threonine that are phosphorylated *in vivo*. HORMAD1 fragment 17-239 contains the full length of HORMA domain (amino acid positions 25-227). n. t.=not tested.

**Table S2.**
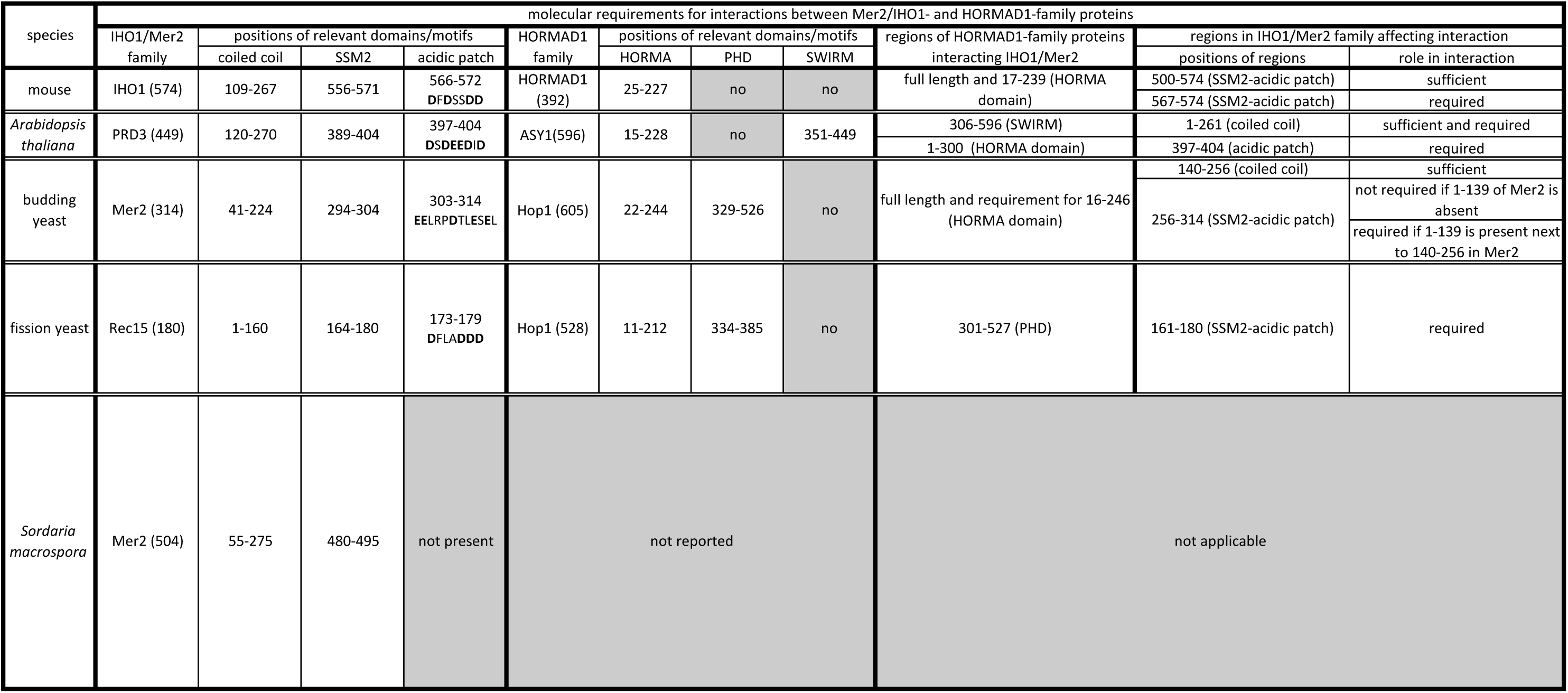
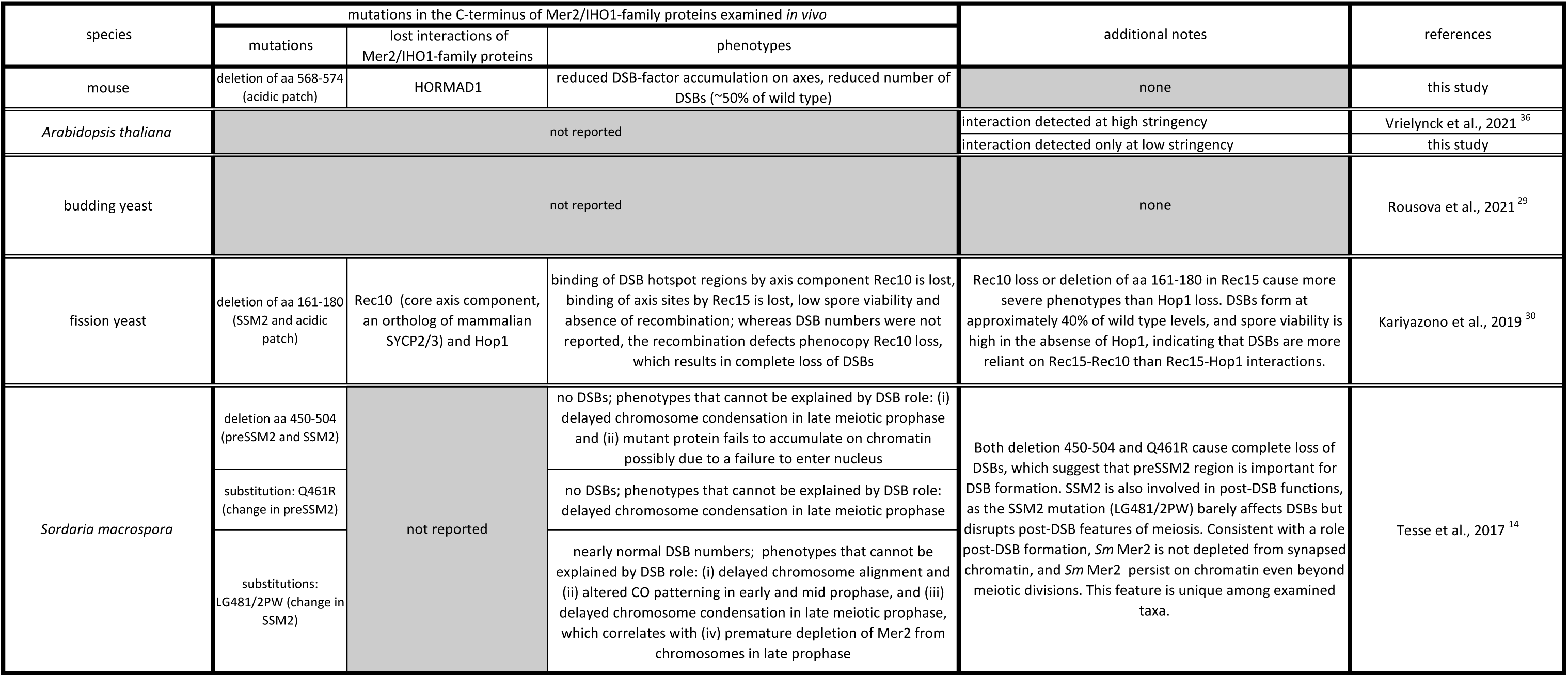
Molecular features underlying the interactions between Mer2/IHO1-and HORMAD1-family proteins and the reported roles of the C-terminal region of Mer2/IHO1-family proteins in diverse taxa (relevant to Fig. 1) The lengths of proteins are indicated in brackets, and amino acid (aa) positions are indicated for motifs/domains, interacting regions and mutations. Amino acid positions for predicted domains/motifs are shown according to publications in the cases of (i) coiled coil in IHO1/Mer2 family ^73^, (ii) SSM2 in IHO1/Mer2 family ^14^, (iii) acidic patch in IHO1/Mer2 family (this study), (iv) HORMA domain and PHD domain in budding yeast ^106^ or according to Uniprot database in the cases of HORMA, PHD and SWIRM domains in mouse, *Arabidopsis thaliana*, and fission yeast. If the table indicates that a region is required for interaction, the requirement was tested with otherwise full-length proteins.

**Table S3.** MS analysis of *in vitro* phosphorylated C-terminal peptides of IHO1 (relevant to Fig. 2) Calculated fragments of NLLCDPDFDS(pS)DNF-COOH and –NH2 peptides corresponding to the MS/MS spectra shown in Figure 2D and S2D, respectively. Identified b- and y-ions are shown in red and blue, respectively. b- and y-ions with neutral losses of phosphorylation (bold), H_2_O and NH_3_ are also indicated in the table.

| #1 | b<sup>-</sup>-Phos | b<sup>2+</sup>-Phos | b<sup>+</sup> | b<sup>2+</sup> | b<sup>3+</sup> | Seq. | y<sup>+</sup> | y<sup>2+</sup> | y<sup>3+</sup> | y<sup>-</sup>-Phos | y<sup>2-</sup>-Phos | #2 |
| --- | --- | --- | --- | --- | --- | --- | --- | --- | --- | --- | --- | --- |
| 1 |  |  | 115.05020 | 58.02874 | 39.02159 | N |  |  |  |  |  | 15 |
| 2 |  |  | **228.13427** | 114.57077 | 76.71627 | L | 1682.59277 | 841.80003 | **561.53578** | 1584.61588 | 792.81158 | 14 |
| 3 |  |  | **341.21833** | 171.11280 | 114.41096 | L | 1569.50871 | **785.25799** | 523.84109 | 1471.53182 | **736.26955** | 13 |
| 4 |  |  | **444.22752** | 222.61740 | 148.74736 | C | 1456.42465 | **728.71596** | 486.14640 | 1358.44775 | **679.72751** | 12 |
| 5 |  |  | **559.25446** | **280.13087** | 187.08967 | D | 1353.41546 | 677.21137 | 451.81001 | 1255.43857 | 628.22292 | 11 |
| 6 |  |  | **656.30722** | 328.65725 | 219.44059 | P | 1238.38852 | **619.69790** | 413.46769 | 1140.41162 | **570.70945** | 10 |
| 7 |  |  | **771.33417** | **386.17072** | 257.78291 | D | **1141.33576** | **571.17152** | 381.11677 | 1043.35886 | 522.18307 | 9 |
| 8 |  |  | **918.40258** | **459.70493** | 306.80571 | F | **1026.30881** | **513.65804** | 342.77446 | 928.33192 | **464.66960** | 8 |
| 9 |  |  | **1033.42952** | **517.21840** | 345.14803 | D | **879.24040** | **440.12384** | 293.75165 | **781.26350** | **391.13539** | 7 |
| 10 |  |  | 1120.46155 | **560.73441** | 374.15870 | S | **764.21346** | **382.61037** | 255.40934 | **666.23656** | 333.62192 | 6 |
| 11 | 1189.48302 | 595.24515 | 1287.45991 | 644.23359 | 429.82482 | S-Phospho | **677.18143** | 339.09435 | 226.39866 | 579.20453 | 290.10590 | 5 |
| 12 | 1304.50996 | **652.75862** | 1402.48685 | **701.74707** | 468.16714 | D | **510.18307** | 255.59517 | 170.73254 |  |  | 4 |
| 13 | 1419.53690 | **710.27209** | 1517.51380 | **759.26054** | 506.50945 | D | **395.15613** | 198.08170 | 132.39023 |  |  | 3 |
| 14 | 1533.57983 | **767.29355** | 1631.55672 | **816.28200** | 544.52376 | N | **280.12918** | 140.56823 | 94.04791 |  |  | 2 |
| 15 |  |  |  |  |  | F | 166.08626 | 83.54677 | 56.03360 |  |  | 1 |

Fragments with neutral losses of H<sub>2</sub>O and NH<sub>3</sub>
| #1 | b-H<sub>2</sub>O<sup>+</sup> | b-H<sub>2</sub>O<sup>2+</sup> | b-H<sub>2</sub>O<sup>3+</sup> | b-NH<sub>3</sub><sup>+</sup> | b-NH<sub>3</sub><sup>2+</sup> | b-NH<sub>3</sub><sup>3+</sup> | Seq. | y-H<sub>2</sub>O<sup>+</sup> | y-H<sub>2</sub>O<sup>2+</sup> | y-H<sub>2</sub>O<sup>3+</sup> | y-NH<sub>3</sub><sup>+</sup> | y-NH<sub>3</sub><sup>2+</sup> | y-NH<sub>3</sub><sup>3+</sup> | #2 |
| --- | --- | --- | --- | --- | --- | --- | --- | --- | --- | --- | --- | --- | --- | --- |
| 1 |  |  |  | 98.02365 | 49.51547 | 33.34607 | N |  |  |  |  |  |  | 15 |
| 2 |  |  |  | 211.10772 | 106.05750 | 71.04076 | L | 1664.58221 | 832.79474 | **555.53225** | 1665.56623 | 833.28675 | **555.86026** | 14 |
| 3 |  |  |  | 324.19178 | 162.59953 | 108.73545 | L | 1551.49815 | **776.25271** | 517.83757 | 1552.48216 | **776.74472** | 518.16557 | 13 |
| 4 |  |  |  | 427.20097 | 214.10412 | 143.07184 | C | 1438.41408 | **719.71068** | 480.14288 | 1439.39810 | **720.20269** | 480.47088 | 12 |
| 5 | **541.24389** | 271.12559 | 181.08615 | **542.22791** | 271.61759 | 181.41415 | D | 1335.40490 | 668.20609 | 445.80648 | 1336.38891 | **668.69810** | 446.13449 | 11 |
| 6 | 638.29666 | 319.65197 | 213.43707 | 639.28067 | 320.14398 | 213.76508 | P | 1220.37796 | **610.69262** | 407.46417 | 1221.36197 | **611.18462** | 407.79217 | 10 |
| 7 | 753.32360 | 377.16544 | 251.77938 | 754.30762 | 377.65745 | 252.10739 | D | 1123.32519 | 562.16623 | 375.11325 | 1124.30921 | 562.65824 | 375.44125 | 9 |
| 8 | 900.39202 | 450.69965 | 300.80219 | **901.37603** | **451.19165** | 301.13019 | F | **1008.29825** | **504.65276** | 336.77093 | **1009.28226** | **505.14477** | 337.09894 | 8 |
| 9 | **1015.41896** | **508.21312** | 339.14450 | **1016.40297** | **508.70513** | 339.47251 | D | **861.22983** | **431.11856** | 287.74813 | **862.21385** | **431.61056** | 288.07613 | 7 |
| 10 | 1102.45099 | **551.72913** | 368.15518 | 1103.43500 | **552.22114** | 368.48319 | S | **746.20289** | 373.60508 | 249.40581 | **747.18691** | **374.09709** | 249.73382 | 6 |
| 11 | 1269.44935 | 635.22831 | 423.82130 | 1270.43336 | 635.72032 | **424.14930** | S-Phospho | 659.17086 | 330.08907 | 220.39514 | 660.15488 | 330.58108 | 220.72314 | 5 |
| 12 | 1384.47629 | 692.74178 | 462.16361 | 1385.46030 | **693.23379** | 462.49162 | D | 492.17250 | 246.58989 | 164.72902 | 493.15652 | 247.08190 | 165.05702 | 4 |
| 13 | 1499.50323 | **750.25525** | 500.50593 | 1500.48725 | **750.74726** | 500.83393 | D | 377.14556 | 189.07642 | 126.38670 | **378.12958** | 189.56843 | 126.71471 | 3 |
| 14 | 1613.54616 | **807.27672** | 538.52024 | 1614.53017 | **807.76873** | **538.84824** | N |  |  |  | **263.10263** | 132.05496 | 88.37240 | 2 |
| 15 |  |  |  |  |  |  | F |  |  |  |  |  |  | 1 |

NLLCDPDFDS(pS)DDNF-NH<sub>2</sub> corresponding to Fig. S2D Fragments and Phosphorylation losses (yellow)
| #1 | b<sup>-</sup>-Phos | b<sup>2+</sup>-Phos | b<sup>+</sup> | b<sup>2+</sup> | b<sup>3+</sup> | Seq. | y<sup>+</sup> | y<sup>2+</sup> | y<sup>3+</sup> | y<sup>-</sup>-Phos | y<sup>2-</sup>-Phos | #2 |
| --- | --- | --- | --- | --- | --- | --- | --- | --- | --- | --- | --- | --- |
| 1 |  |  | 115.05020 | 58.02874 | 39.02159 | N |  |  |  |  |  | 15 |
| 2 |  |  | **228.13427** | 114.57077 | 76.71627 | L | 1681.60876 | 841.30802 | **561.20777** | 1583.63186 | 792.31957 | 14 |
| 3 |  |  | **341.21833** | 171.11280 | 114.41096 | L | 1568.52470 | **784.76599** | 523.51308 | 1470.54780 | **735.77754** | 13 |
| 4 |  |  | **444.22752** | 222.61740 | 148.74736 | C | 1455.44063 | **728.22395** | 485.81839 | 1357.46374 | 679.23551 | 12 |
| 5 |  |  | **559.25446** | 280.13087 | 187.08967 | D | 1352.43145 | 676.71936 | 451.48200 | 1254.45455 | 627.73091 | 11 |
| 6 |  |  | **656.30722** | 328.65725 | 219.44059 | P | 1237.40450 | **619.20589** | 413.13969 | 1139.42761 | 570.21744 | 10 |
| 7 |  |  | **771.33417** | **386.17072** | 257.78291 | D | 1140.35174 | 570.67951 | 380.78876 | 1042.37484 | 521.69106 | 9 |
| 8 |  |  | **918.40258** | **459.70493** | 306.80571 | F | **1025.32480** | **513.16604** | 342.44645 | 927.34790 | 464.17759 | 8 |
| 9 |  |  | **1033.42952** | **517.21840** | 345.14803 | D | **878.25638** | **439.63183** | 293.42365 | 780.27949 | 390.64338 | 7 |
| 10 |  |  | 1120.46155 | **560.73441** | 374.15870 | S | **763.22944** | **382.11836** | 255.08133 | **665.25255** | 333.12991 | 6 |
| 11 | 1189.48302 | 595.24515 | 1287.45991 | **644.23359** | 429.82482 | S-Phospho | **676.19741** | 338.60234 | 226.07066 | 578.22052 | 289.61390 | 5 |
| 12 | 1304.50996 | **652.75862** | 1402.48685 | **701.74707** | 468.16714 | D | **509.19905** | 255.10316 | 170.40454 |  |  | 4 |
| 13 | 1419.53690 | **710.27209** | 1517.51380 | 759.26054 | 506.50945 | D | **394.17211** | 197.58969 | 132.06222 |  |  | 3 |
| 14 | 1533.57983 | **767.29355** | 1631.55672 | **816.28200** | **544.52376** | N | 279.14517 | 140.07622 | 93.71991 |  |  | 2 |
| 15 |  |  |  |  |  | F-Amidated | 165.10224 | 83.05476 | 55.70560 |  |  | 1 |

| #1 | b-H<sub>2</sub>O<sup>+</sup> | b-H<sub>2</sub>O<sup>2+</sup> | b-H<sub>2</sub>O<sup>3+</sup> | b-NH<sub>3</sub><sup>+</sup> | b-NH<sub>3</sub><sup>2+</sup> | b-NH<sub>3</sub><sup>3+</sup> | Seq. | y-H<sub>2</sub>O<sup>+</sup> | y-H<sub>2</sub>O<sup>2+</sup> | y-H<sub>2</sub>O<sup>3+</sup> | y-NH<sub>3</sub><sup>+</sup> | y-NH<sub>3</sub><sup>2+</sup> | y-NH<sub>3</sub><sup>3+</sup> | #2 |
| --- | --- | --- | --- | --- | --- | --- | --- | --- | --- | --- | --- | --- | --- | --- |
| 1 |  |  |  | 98.02365 | 49.51547 | 33.34607 | N |  |  |  |  |  |  | 15 |
| 2 |  |  |  | 211.10772 | 106.05750 | 71.04076 | L | 1663.59819 | 832.30274 | **555.20425** | 1664.58221 | 832.79474 | **555.53225** | 14 |
| 3 |  |  |  | 324.19178 | 162.59953 | 108.73545 | L | 1550.51413 | **775.76070** | 517.50956 | 1551.49815 | **776.25271** | 517.83757 | 13 |
| 4 |  |  |  | 427.20097 | 214.10412 | 143.07184 | C | 1437.43007 | **719.21867** | 479.81487 | 1438.41408 | **719.71068** | 480.14288 | 12 |
| 5 | **541.24389** | 271.12559 | 181.08615 | 542.22791 | 271.61759 | 181.41415 | D | 1334.42088 | 667.71408 | 445.47848 | 1335.40490 | **668.20609** | 445.80648 | 11 |
| 6 | 638.29666 | 319.65197 | 213.43707 | 639.28067 | 320.14398 | 213.76508 | P | 1219.39394 | 610.20061 | 407.13616 | 1220.37795 | **610.69262** | 407.46417 | 10 |
| 7 | 753.32360 | 377.16544 | 251.77938 | 754.30762 | 377.65745 | 252.10739 | D | 1122.34118 | 561.67423 | 374.78524 | 1123.32519 | 562.16623 | 375.11325 | 9 |
| 8 | 900.39202 | 450.69965 | 300.80219 | **901.37603** | 451.19165 | 301.13019 | F | 1007.31423 | **504.16075** | 336.44293 | **1008.29825** | **504.65276** | 336.77093 | 8 |
| 9 | **1015.41896** | **508.21312** | 339.14450 | 1016.40297 | **508.70513** | 339.47251 | D | **860.24582** | **430.62655** | 287.42012 | **861.22983** | **431.11856** | 287.74813 | 7 |
| 10 | 1102.45099 | **551.72913** | 368.15518 | 1103.43500 | **552.22114** | 368.48319 | S | **745.21888** | 373.11308 | 249.07781 | **746.20289** | **373.60508** | 249.40581 | 6 |
| 11 | 1269.44935 | 635.22831 | 423.82130 | 1270.43336 | 635.72032 | **424.14930** | S-Phospho | 658.18685 | 329.59706 | 220.06713 | **659.17086** | 330.08907 | 220.39514 | 5 |
| 12 | 1384.47629 | 692.74178 | 462.16361 | 1385.46030 | **693.23379** | 462.49162 | D | 491.18849 | 246.09788 | 164.40101 | **492.17250** | 246.58989 | 164.72902 | 4 |
| 13 | 1499.50323 | **750.25525** | 500.50593 | 1500.48725 | 750.74726 | 500.83393 | D | 376.16154 | 188.58441 | 126.05870 | 377.14556 | 189.07642 | 126.38670 | 3 |
| 14 | 1613.54616 | **807.27672** | **538.52024** | 1614.53017 | **807.76873** | **538.84824** | N |  |  |  | **262.11862** | 131.56295 | 88.04439 | 2 |
| 15 |  |  |  |  |  |  | F-Amidated |  |  |  |  |  |  | 1 |

**Table S4.** Fertility checks in *Iho1^C7Δ/C7Δ^* mice (relevant to Fig. 3) Quantification of pup numbers from crosses of wild type to wild type (wt) and wild type to *Iho1^C7Δ/C7Δ^* mice. Statistical significance was calculated by two tailed t-test with Welch correction.

| female age |  |  | female < 28 weeks |  | female >28 weeks |  |
| --- | --- | --- | --- | --- | --- | --- |
| female | wt |  | wt | *lho1*<sup>C7Δ/C7Δ</sup> | wt | *lho1*<sup>C7Δ/C7Δ</sup> |
| male | wt | *lho1*<sup>C7Δ/C7Δ</sup> | wt |  |  |  |
| breeding pairs | 5 | 5 | 9 | 12 | 9 | 12 |
| pups/breeding week | 1,231 | 1,426 | 1,331 | 1,205 | 1,168 | 0,8285 |
| total breeding weeks | 378 | 416 | 175 | 234 | 151 | 206 |
| Significance, P= | 0,3584 |  | 0,1356 |  | 0,0106 |  |

**Table S5.**
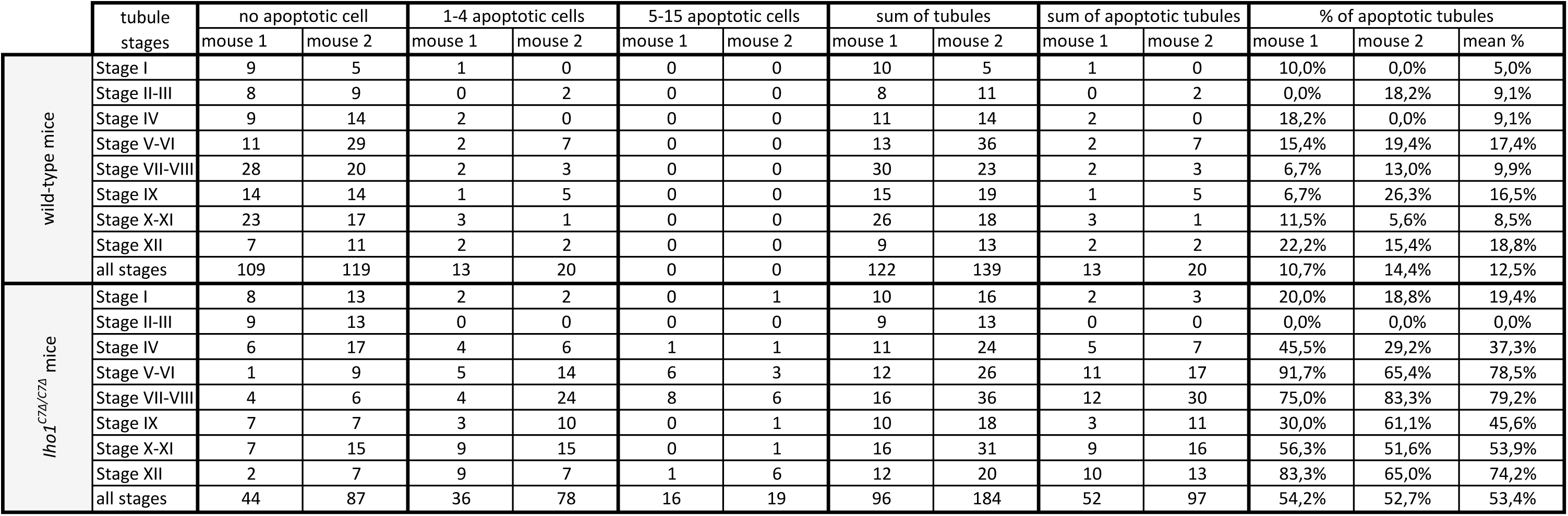
Spermatocyte apoptosis in Iho1^C7Δ/C7Δ^ mice (relevant to Fig. 3) Quantification of seminiferous tubules where apoptosis was detected in the cell layer consisting of pachytene, diplotene or meiotically dividing cells. Stages of the epithelial seminiferous cycle are indicated. Apoptosis was detected by immunostaining for cleaved PARP.

**Table S6.**
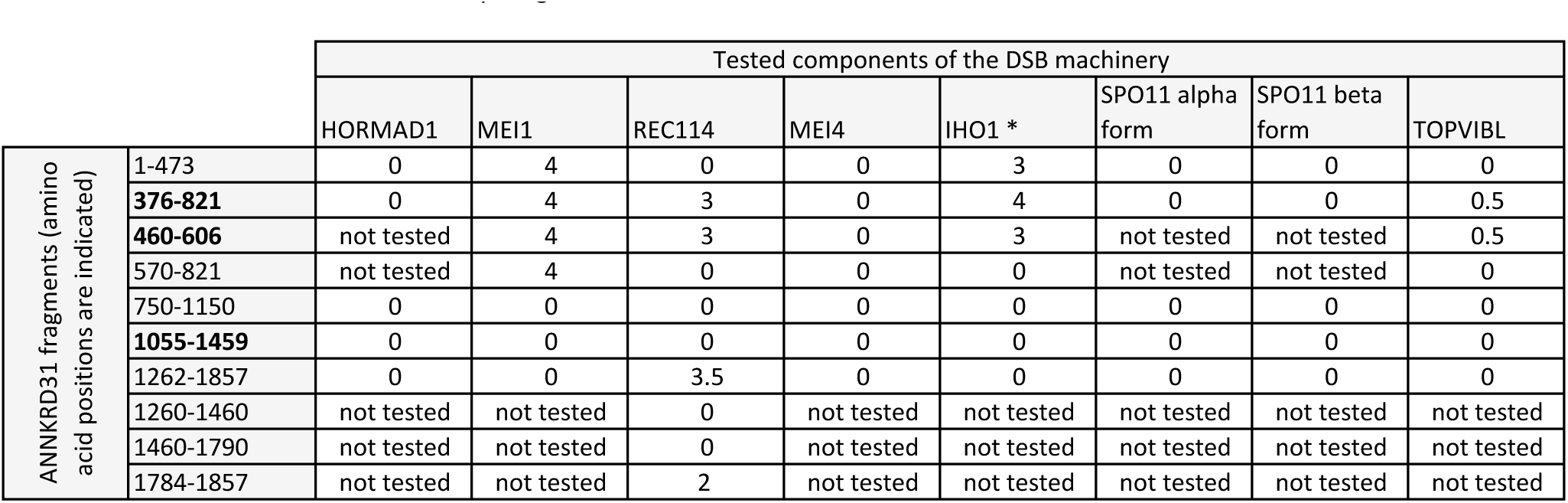
Y2H interactions between ANKRD31 fragments (Gal4 AD) and key components of the DSB machinery (Gal4 BD) (relevant to Fig. 7) Growth of budding yeast in Y2H assay (-Leu, -Trp, -His, -Ade plates) was graded between 0 (no growth) and 4 (very strong growth). Bold fragments 376-821 and 460-606 contained ankyrin repeats 1-3 (amino acid positions 475-570) and fragment 1055-1459 contained ankyrin repeats 4-6 (amino acid positions 1162-1257). * IHO1-GAL4BD fusion mildly autoactivates leading to weak growth beyond three days, hence IHO1 interactions were scored after two days of growth on selective media. n. t.=not tested.

